# Size-dependent secondary nucleation and amplification of α-synuclein amyloid fibrils

**DOI:** 10.1101/2021.12.28.474324

**Authors:** Arunima Sakunthala, Debalina Datta, Ambuja Navalkar, Laxmikant Gadhe, Pradeep Kadu, Komal Patel, Surabhi Mehra, Rakesh Kumar, Debdeep Chatterjee, Kundan Sengupta, Ranjith Padinhateeri, Samir K. Maji

**Author notes:** Corresponding author: Samir K. Maji, Department of Biosciences and Bioengineering, IIT Bombay, Powai, Mumbai 400 076, India.

## Abstract

The size of the amyloid seeds is known to modulate their autocatalytic amplification and cellular toxicity. However, the seed size-dependent secondary nucleation mechanism, toxicity, and disease-associated biological processes mediated by α-synuclein (α-Syn) fibrils are largely unknown. Using the cellular model and *in vitro* reconstitution, we showed that the size of α-Syn fibril seeds not only dictates its cellular internalization and associated cell death; but also the distinct mechanisms of fibril amplification pathways involved in the pathological conformational change of α-Syn. Specifically, small-sized fibril seeds showed elongation possibly through monomer addition at the fibril termini; whereas longer fibrils template the fibril amplification by surface-mediated nucleation as demonstrated by super-resolution microscopy. The distinct mechanism of fibril amplification, and cellular uptake along with toxicity suggest that breakage of fibrils into different sizes of seeds determine the underlying pathological outcome of synucleinopathies.

## Introduction

Misfolding of α-synuclein (α-Syn) and intraneuronal accumulation of amyloid aggregates are the key neuropathological features associated with synucleinopathies including Parkinson’s disease (PD) (Spillantini et al., 1997, Spillantini and Goedert, 2000). Although fibrillization is a prerequisite for Lewy body (LB) formation (Goedert et al., 2017, Lashuel, 2020), the precise cellular and molecular mechanism, which drives α-Syn aggregation and subsequent disease progression are still unclear (Mahul-Mellier et al., 2020). Multiple evidence suggests that the rate of α-Syn aggregation and its propagation critically determine the progression of PD (Gómez-Benito et al., 2020). Intriguingly, the mechanism of α-Syn aggregation is a nucleation-dependent polymerization phenomenon (Wood et al., 1999), wherein the natively unstructured α-Syn monomers undergo self-assembly and lead to the formation of thermodynamically stable amyloid aggregates (Chiti and Dobson, 2006) through various microscopic processes (Cohen et al., 2012, Buell et al., 2014). The rate of formation of these assemblies can be regulated by dominant secondary processes including fibril fragmentation and seeded-nucleation events (Xue et al., 2008, Knowles et al., 2009, Arosio et al., 2015, Marrero-Winkens et al., 2020). For instance; in the seeded aggregation model, the assembly of amyloid fibrils occurs either through templated addition of monomers on the growth competent ends of preformed fibrils (PFF) via fibril elongation (Collins et al., 2004, Ferrone, 1999, Xue, 2015, Pinotsi et al., 2014), or interaction/binding of monomers on the catalytic surface of fibril species by surface-mediated secondary nucleation (Zimmermann et al., 2021, Gaspar et al., 2017, Kumari et al., 2021). Of note, these microscopic processes rely on multiple parameters including native protein conformations and other external factors, which may co-occur and generate amyloid aggregates with distinct pathological phenotypes (Buell et al., 2014, Linse, 2019, Koloteva-Levine et al., 2021).

Interestingly, fibril fragmentation, which leads to the formation of shorter fibril seeds is demonstrated as a vital step for the amplification of protein aggregates and the spreading of prion seeds (Aguzzi and Rajendran, 2009, Xue et al., 2010, Meisl et al., 2020). This may accelerate the transmission of pathological inclusions into various regions of the brain (Luk et al., 2009). However, under normal conditions, protein homeostasis is tightly regulated by protein folding quality control machinery, which maintains the proteome integrity and limits the accumulation of protein aggregates (Dobson, 1999, Lansbury, 1999). Hence, fragmentation can be considered an inherent biological property of amyloid fibrils, which is typically modulated by numerous factors including thermal motion, shear forces, direct mechanical stress, and catalytic activity of molecular chaperones in cells (Xue et al., 2009, Beal et al., 2020, Shorter and Lindquist, 2004). For example; fibril disassembly by the Human heat shock protein 70 (HSP 70) family promotes the disaggregation of amyloids (such as α-Syn and Tau) into potent species with high seeding ability and prion-like behaviour (Winkler et al., 2012, Tittelmeier et al., 2020, Nachman et al., 2020). Remarkably, a growing body of evidence suggests that biological attributes of amyloid fibrils can be regulated by the size of the fibrils, which is governed by the magnitude of the fragmentation events to a greater extent (Xue et al., 2008, Xue et al., 2010, Tanaka et al., 2006). In this context, extensive research has been conducted to understand the correlation between the fragmentation of amyloid fibrils and their cytotoxic potential (Aguzzi and Lakkaraju, 2016, Tarutani et al., 2016, Marchante et al., 2017). However, the interplay between the heterogeneity in α-Syn fibril size and the mechanism of seed size-dependent secondary nucleation pathways involved in the prion-like transmission of pathological conformers is elusive. Here, we hypothesize that apart from biological/pathological features, the heterogeneity and nanoscale differences in fibril size may regulate the progression of α-Syn pathology by dictating distinct secondary nucleation mechanisms associated with fibril amplification pathways.

To test this, we generated α-Syn fibril fragments of variable length by controlled time-dependent sonication. We systematically examined the size-dependent effects mediated by α-Syn fibril seeds in PD pathogenesis by probing the mechanistic events involved in amyloid amplification and disease-associated cellular processes using a combination of biophysical and cell-based studies. The controlled fragmentation protocol generates α-Syn fibril fragments of variable lengths but with identical cross-β-sheet structures. When added exogenously, the internalization behaviour and cellular seeding potential of these fibril fragments vary with seed length. The seed length also modulates the number and size of seeding-dependent α-Syn aggregates generated in cells overexpressing α-Syn. In addition, extensive global analysis of *in vitro* aggregation kinetics together with dual-colour super-resolution microscopy provided a direct demonstration that short fibril seeds promote fibril amplification by templated elongation, whereas long fibril seeds favours fibril amplification by surface-mediated secondary nucleation. Further, the short α-Syn fibril fragments with higher uptake efficiency showed enhanced cellular toxicity, higher membrane damage potential, and cellular apoptosis compared to longer fibril fragments. Overall, the present study provides the fact that fragmentation of fibrils and resulting nanoscale differences in α-Syn amyloid seeds might regulate the critical cellular phenomena associated with PD pathogenesis including fibril amplification and prion-like spreading of the disease pathology.

## Results

### Generation and biophysical characterization of α-Syn fibril seeds

It has been established that fibril fragmentation is critical for the propagation of misfolded prion proteins (Pezza and Serio, 2007, Kundel et al., 2018, Scheckel and Aguzzi, 2018, Marrero-Winkens et al., 2020). To explore, how the fragmentation-dependent seed length of α-Syn fibrils determines the amyloid amplification mechanism and disease-associated cellular activities, we prepared α-Syn fibrils by incubation under physiological conditions and rationally generated different-sized fibril fragments by controlled time-dependent sonication (Figures S1). The morphological analysis by transmission electron microscopy (TEM) and atomic force microscopy (AFM) confirmed a significant difference in the length profile among the fibril fragments prepared by time-dependent sonication. We found the typical long unbranched fibril morphology for unsonicated fibril samples; whereas short fibril fragments were observed after sonication. The length of the fibril fragments decreased with an increase in the extent of sonication time (Figure 1A-C). The trend in the particle size distribution of the fragmented fibrils observed by dynamic light scattering (DLS) was also consistent with the fibril length distribution profile obtained from TEM image analysis (Figure 1D). However, the secondary structural analysis of fibril fragments monitored by circular dichroism (CD) (Figure S2D) and Fourier transform infrared (FTIR) spectroscopy suggests that the structure of α-Syn amyloid fibrils remains unchanged during the fragmentation process (Figure 1E, S2E). This was further supported by a similar extent of Thioflavin T (ThT) fluorescence (Figure S2A) and a unique cross-β-diffraction pattern shown by the different-sized fibril seeds (Figure 1F). We further examined the possibility that sonication and subsequent fragmentation might alter the exposed hydrophobic surfaces, which may dictate their physicochemical and biological activities. Interestingly, the ANS and NR fluorescence (the dyes bind to the exposed hydrophobic surfaces of the protein) study showed that fragmentation did not affect the binding and fluorescence of both the dyes suggesting no alterations in the exposed hydrophobic surfaces (Figure S2B, C). Moreover, SDS-PAGE analysis showed a single band at ∼17 kDa indicating no signs of protein degradation during fragmentation of α-Syn fibrils (Figure S2F). Overall, the biophysical characterization of fragmented fibrils indicates that fragmentation did not change the characteristic secondary structural conformation of α-Syn amyloid fibrils; but it critically affected the physical attributes of fibrils, which is predominantly reflected in its length.

**Figure 1:**
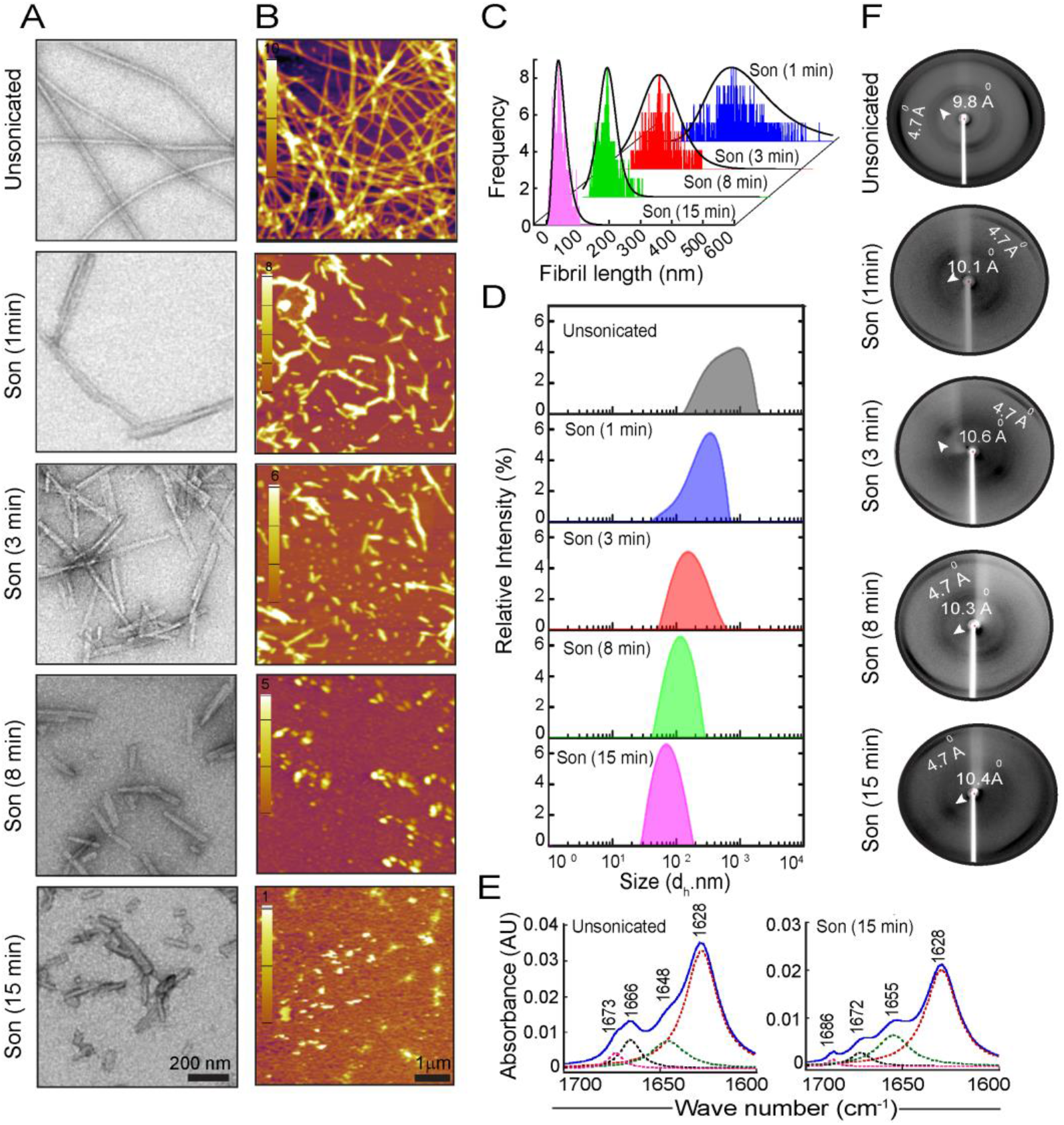
Morphological and biophysical characterization of α-Syn fibril fragments. Fragmentation of α-Syn fibrils was done by controlled time-dependent sonication (1 min, 3 min, 8 min, and 15 min); unfragmented fibrils (denoted as unsonicated) were used as a control throughout the study. (A) TEM images and (B) AFM images showing the morphological differences in the fibril fragments prepared by time-dependent sonication. The scale bar is 200 nm for TEM images and 1 μm for AFM images. (C) The curve fitted 3D plot showing the length distribution of the fibril fragments calculated from TEM images using ImageJ analysis (n>260 counts). Fragmentation of the fibrils significantly decreased the fibril length; the length distribution profile varies with the extent of sonication. (D) Size distribution analysis of α-Syn fibril fragments by DLS showing a significant decrease in the size of the fibril fragments (represented as d_h_. nm) with an increase in the extent of sonication. (E) The curve-fitted FTIR spectra (1600-1700 cm^-1^) together with Fourier deconvolution showing no changes in the secondary structure after extended sonication in comparison to unsonicated fibrils. (F) The XRD analysis of α-Syn fibril fragments of variable lengths showing the characteristics of cross-β-sheet reflections (meridional arcs at 4.7 Å and equatorial arcs between ∼8-11 Å), indicating that the core secondary structure of amyloids remains unchanged during the fragmentation of fibrils *in vitro*.

### Size-dependent internalization and templating behaviour of α-Syn fibrils with variable lengths

It has been demonstrated that seed-dependent misfolding of native proteins and the spontaneous formation and propagation of their amyloid state are fundamental cellular events in the spreading of various prion-like disorders including PD (Krammer et al., 2009, Frost et al., 2009, Jucker and Walker, 2018). In this regard, various studies indicated that cellular uptake, templating/seeding, and intercellular transmission of α-Syn fibrils are the key events contributing to the onset and the propagation of LB pathology in PD brains (Steiner et al., 2011, Hansen et al., 2011, Kordower et al., 2008). Since fragmentation events increase the availability of fibril termini (Linse, 2019), it may facilitate its direct binding with the plasma membrane (Xue et al., 2010), and consequently aggravate the cellular internalization process. To examine the seed-size dependent cellular uptake of α-Syn fibrils, we performed internalization studies using exogenous addition of (50 nM) fragmented α-Syn fibrils seeds of variable lengths in SH-SY5Y human neuroblastoma cells. Prior optimization studies were done using α-Syn fibrils seeds (using an intermediate seed length; Son (3 min)), which suggest that very low concentration of fibril seeds (50 nM) can be successfully used for the internalization experiments without the interference from the high intensity of fluorescence signal (Figure S3 A-C). Notably, the confocal microscopy imaging suggested that fibril uptake in cells was in inverse correlation with fibril seed length (Figure 2A; S4). The quantification of the amount of internalized fibrils was done using FACS analysis, which is well corroborated with the respective confocal microscopy imaging data (Figure 2B, C). Interestingly, a similar observation was also found in differentiated SH-SY5Y (SH-SY5Y (D)) cells (Figure S5). Moreover, the short α-Syn fibril fragments internalized more rapidly and showed the highest cellular uptake within shorter incubation time (3h) as compared to the longer fibril seeds (Figure S3D), suggesting that short fibril fragments might possess a higher potential to initiate the disease pathogenesis by accelerating the cellular uptake of misfolded seeds into the cells.

**Figure 2:**
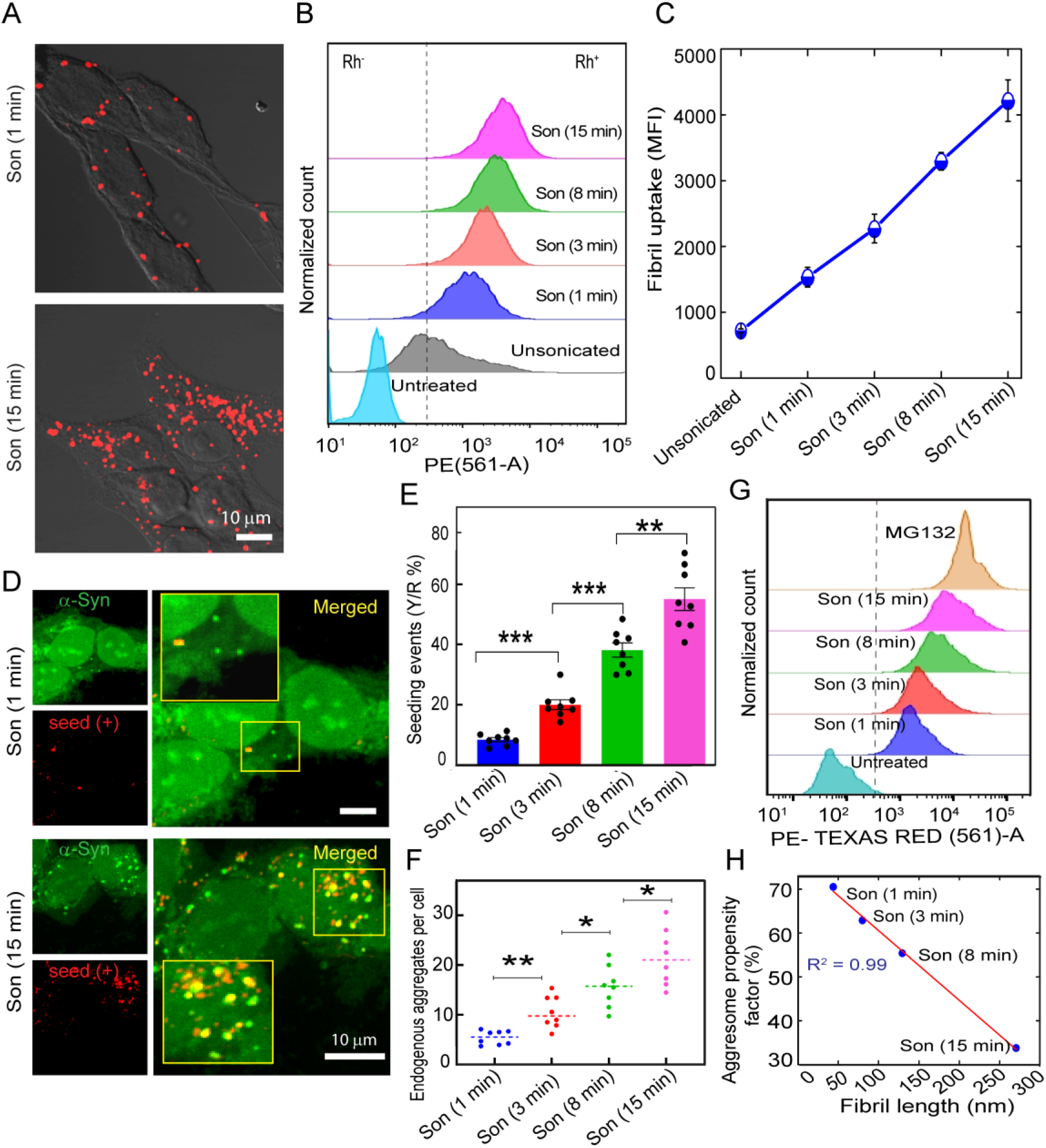
Size-dependent internalization and templating behaviour of α-Syn fibrils seeds with variable lengths. (A) Confocal laser scanning imaging of SH-SY5Y cells treated with Rhodamine (Rh)-labelled α-Syn fibril fragments (50 nM) of varying lengths showing higher internalization by short fibril seeds compared to longer seeds. SH-SY5Y cells treated with unsonicated α-Syn fibrils were used as control. (B) The FACS analysis and (C) corresponding median fluorescence intensity (MFI) values represent the internalization efficiency of α-Syn fibril fragments in a size-dependent manner. Cells treated with unsonicated fibrils were used as control. (D) Confocal images of SH-SY5Y cells stably expressing C4 tagged-α-Syn showing seeding events and aggregate formation by the endogenous protein due to exogenous addition of 50 nM of Rh-labelled Syn fibril seeds of variable lengths. Internalized α-Syn fibril seeds were observed as red punctate structures. The merged images from each group were magnified to highlight seeding events (colocalization, observed as yellow) and amplification of endogenous protein (observed as green inclusions) in cells. The scale bar is 10 μM. (E) Relative seeding events (yellow/red, Y/R ratio) were quantified in cells using Image J from confocal images from three independent sets of experiments with n>80 cells. (F) The number of endogenous protein aggregates (green inclusion) quantified from confocal images of FlAsH stained C4 α-Syn SH-SY5Y cells using Image J analysis (from three independent sets; n>80 cells) showing higher fibril amplification by short fibril seeds compared to longer seeds. (G) The nature of endogenous α-Syn aggregates was quantified using aggresome marker, ProteoStat dye followed by FACS analysis showing higher aggresome formation by shorter seeds compared to the longer ones. Cells treated with MG132, a proteasome inhibitor were used as a positive control. (H). The correlation plot (R^2^= 0.99) showing an inverse correlation of the aggresome propensity factor (determined from ProteoStat assay) with fibril lengths (calculated from TEM image analysis). Values represented as mean ± SEM, n=3 from independent experiments.

We further examined the seeding ability of α-Syn fibril fragments of variable lengths in SH-SY5Y cells stably expressing tetracysteine-tagged α-Syn (C4 α-Syn). This stable cell line has a doxycycline (DOX) inducible system for the controlled expression of α-Syn, which makes it a robust and tunable model for studying seeding and aggregation in cells (Mehra et al., 2020, Ray et al., 2020). The tetracysteine (C4) motif (∼1.5 kDa) binds to biarsenical dyes including FlAsH with high affinity, which enables the visualization and localization of proteins inside the cells (Irtegun et al., 2011, Ray et al., 2020, Whitt and Mire, 2011). Moreover, C4 tagged-α-Syn exhibit equivalent biochemical and biophysical features as of wild-type (WT) α-Syn (Roberti et al., 2007); therefore is an ideal system for studying α-Syn aggregation related to synucleinopathies. Notably, the sensitivity of biarsenical dyes for detecting aggregates is comparable to that of routine antibodies against LBs (Roberti et al., 2007). The fluorescence microscopy studies and subsequent quantification of α-Syn puncta have suggested that compared to unsonicated fibrils, α-Syn fibril fragments have high seeding potential in cells (Figure 2D, S6). Moreover, the highest amount of seeding was observed in cells (∼ 55.10%) treated with shortest fibril fragments (Son (15 min)) followed by Son (8 min) ∼ 38.18% and Son (3 min) fibrils ∼ 20.0%. The least seeding effect (∼ 8.5%) was observed for Son (1 min) fibrils (Figure 2E). This trend of seeding was also correlated with the templating behaviour inside the cells (Figure 2F). Consistent with this, the aggresome marker, ProteoStat binding assay (Shen et al., 2011) also showed enhanced amplification of the endogenous α-Syn aggregates in cells treated with shorter seed lengths (Figure 2G and H). This suggests that short fibril seeds with high seeding potential enhance the aggresome forming potential and amplification of endogenous α-Syn inclusions as compared to longer seeds.

### Distinct intracellular α-Syn inclusion formation depends on the size of the α-Syn fibril seeds

Our results postulate that the extent of seeding and the amount of α-Syn aggregate formation in cells significantly vary depending on the size of the amyloid seeds. Following this, we also observed that the intracellular α-Syn aggregates generated in response to seeding were heterogeneous in size and shape, which also varied with the size of the fibril seeds. To get more insight into this morphological heterogeneity of endogenous α-Syn inclusions, we analysed the α-Syn aggregates generated in C4 tagged**-**α-Syn SH-SY5Y cells by confocal microscopy imaging. Here, three types of cell populations with endogenous aggregates were observed; (i) cells with only spherical aggregates, (ii) cells with a combination of both spherical and filamentous aggregates, and (iii) cells with only filamentous aggregates (Figure 3A). We observed more percentage of cells with elongated filamentous aggregates upon treatment with short α-Syn seeds. On the contrary, longer α-Syn seeds majorly resulted in more cells with spherical inclusions and relatively lesser with elongated aggregates (Figure 3B). Interestingly, the size distributions of the endogenous α-Syn aggregates shifted towards higher values upon seeding with α-Syn short fibrils (Son (15 min)) as compared to the α-Syn aggregates seeded by longer fibril seeds (Son (1 min)) (Figure 3C). It could be possible that, in the case of short fibril seeds, the endogenous α-Syn inclusion formation occurs primarily via templated elongation (monomer addition to fibril end), due to the availability of more growth competent fibril ends (since we used identical concentrations of short and longer seeds). This might result in enhanced fibril amplification. In contrast, fewer elongated aggregates but more spherical inclusions of smaller size were observed in cells that were treated with longer fibril seeds. This might be possible because of the fibril amplification facilitated via surface-mediated secondary nucleation by the longer seeds (Figure 3D). Indeed, fibril surfaces have been reported as heterogeneous nucleation sites for amyloid growth through secondary nucleation pathways (Törnquist et al., 2018, Linse, 2019). Taken together, our results highlight that the size of α-Syn fibril seeds modulates the intracellular seeding potential and endogenous aggregate formation by dictating distinct size-dependent microscopic secondary nucleation mechanisms in cells.

**Figure 3:**
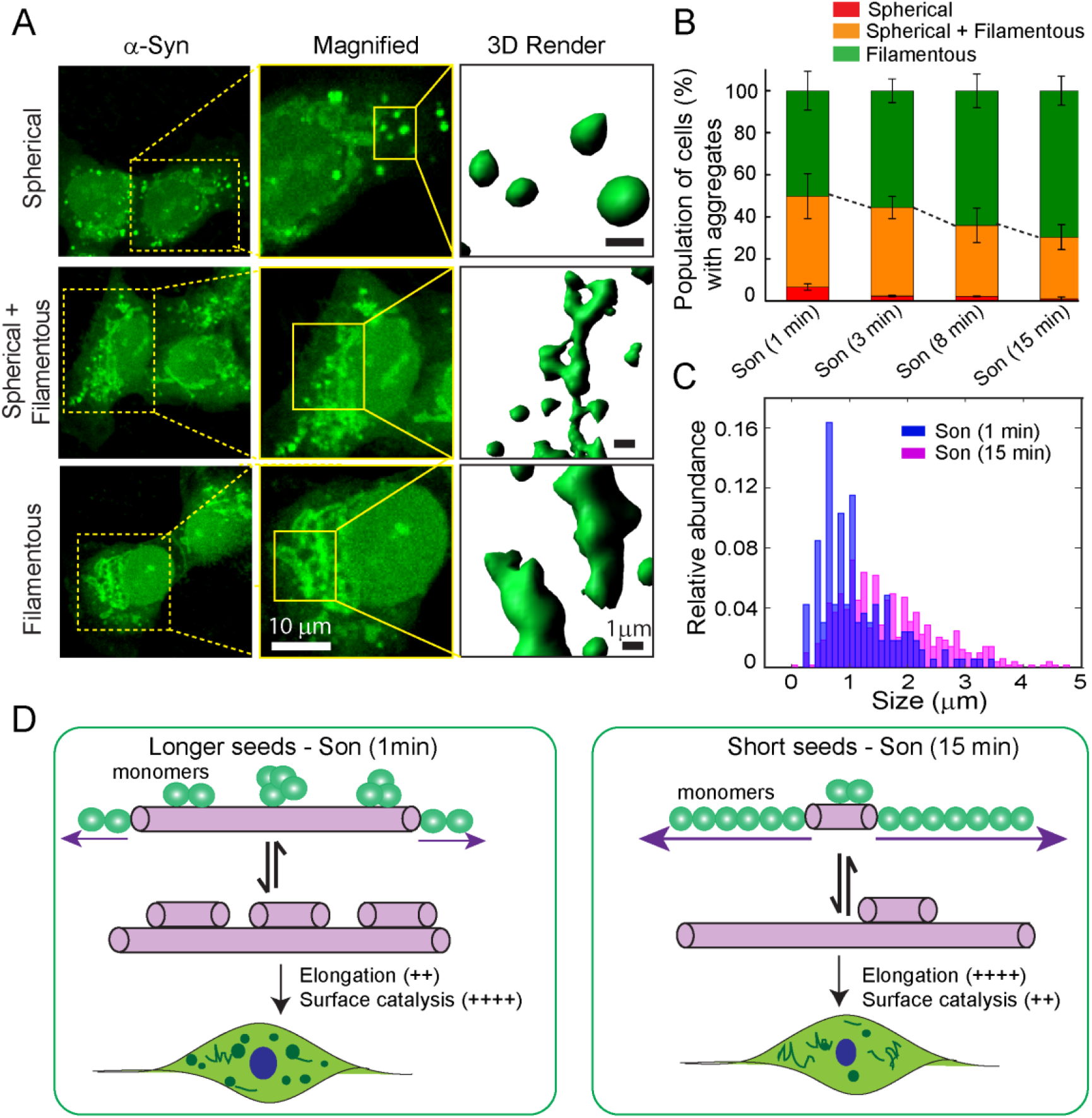
Quantification and morphological characterization of endogenous α-Syn inclusions in C4 tagged-α-Syn SH-SY5Y cells. (A) Confocal images of C4 tagged**-**α-Syn cells showing morphologically distinct endogenous aggregates. Cells with only spherical aggregates (upper panel), cells with both spherical and filamentous aggregates (middle panel), and cells with only elongated filamentous aggregates (lower panel). Rendered representation of the endogenous α-Syn aggregates from confocal Z-stack images using IMARIS showing the distinct morphological characteristics of the intracellular aggregates. The scale bar is 1 μm. (B) Bar plot showing the percentage population of cells with morphologically distinct intracellular aggregates upon seeding with fibrils (50 nM) of variable lengths. The analysis was done from confocal images acquired from two independent sets; cells counted are n > 80. Short fibril seed (Son (15 min)) treatment resulted in a higher percentage of cells with elongated aggregates; whereas treatment with longer fibril seeds (Son (1 min)) showed the lesser percentage of cells with elongated aggregates and a higher percentage with spherical inclusions. Data represented as Mean ± SEM; from 2 independent sets of experiments. (C) Distribution plot for the size of endogenous aggregates generated in cells upon seeding with short (Son (15 min)) and longer fibril seeds (Son (1 min)). Treatment with longer fibril seeds resulted in more aggregates with smaller sizes compared to that with short fibril seeds. The size of the inclusions was analysed from confocal images using Image J software (n=20). (D) Schematic representation illustrating the possible intracellular seeding mechanism exerted by α-Syn fibrils seeds with varying lengths.

### Fibril size dictates the secondary nucleation mechanism of seeded amyloid growth *in vitro*

The kinetic rate of secondary nucleation pathways in amyloid aggregation is highly dependent on factors associated with protein sequences/structure as well as external conditions such as pH, and temperature (Buell et al., 2014, Linse, 2019). We sought to study the extent to which the seed size modulates the primary and secondary nucleation pathways in fibril amplification *in vitro*. The aggregation reaction of monomeric α-Syn (150 μM) was performed in the presence of preformed α-Syn fibril seeds (0.5% v/v) of varying lengths prepared by time-dependent sonication. The aggregation reaction was monitored by ThT fluorescence assay as a function of time. The sigmoidal amyloid growth curve was analyzed using Amylofit, robust web-based software for global analysis of kinetic data (Meisl et al., 2016). The acquired data were fitted to the ‘Secondary Nucleation Dominated’ model following the guideline specifications described by Meisl *et al*. (Meisl et al., 2016) (Table 1). Consistent with the cell data, we also found faster kinetics (shorter lag time) for α-Syn aggregation reaction seeded with short fibril seeds (Son (15 min)) followed by Son (8 min), Son (3 min), and Son (1 min) fibril seeds (Figure 4A-C). Moreover, the seeding ability of these fibrillar seeds is perfectly correlated (R^2^ = 0.99) with its seed length (Figure 4D). Further, we estimated the overall parameters governing primary nucleation (λ) and secondary nucleation (κ) pathways from the estimated values of k_n_ (primary nucleation rate constant), k_+_ (elongation rate constant), and k_2_ (secondary nucleation rate constants) obtained from the fitting (Cohen et al., 2013). A size-dependent increase in elongation rate constant (k_+_) was observed for seeded reaction with fragmented fibrils of decreasing length. The value of λ, (the product of k_n_k_+_, a combined parameter for primary nucleation pathways) was also increased with a decrease in the fibril seed length (Figure 4E). Notably, we observed a higher value for κ, (parameter governing secondary pathway) in seeded reactions with short fibril seeds, suggesting that overall secondary nucleation pathways are more dominant for short fibril seeds relative to their longer counterparts (Figure 4F). Whereas, the elongation rate constant (k_+_) was estimated to be relatively less for longer fibril seeds. This is possibly due to the fact the longer fibril fragments have more exposed surface area, which acts as a catalytic surface for the binding of monomers with high affinity. This would accelerate secondary nucleation via surface-catalyzed secondary nucleation. The calculated values are considered only as estimates to compare the two cases of fibril amplification events. Altogether, the global fit analysis of kinetic data allowed us to estimate the contribution of the different microscopic processes of secondary nucleation in the seed-size-dependent amyloid assembly phenomena.

**Figure 4:**
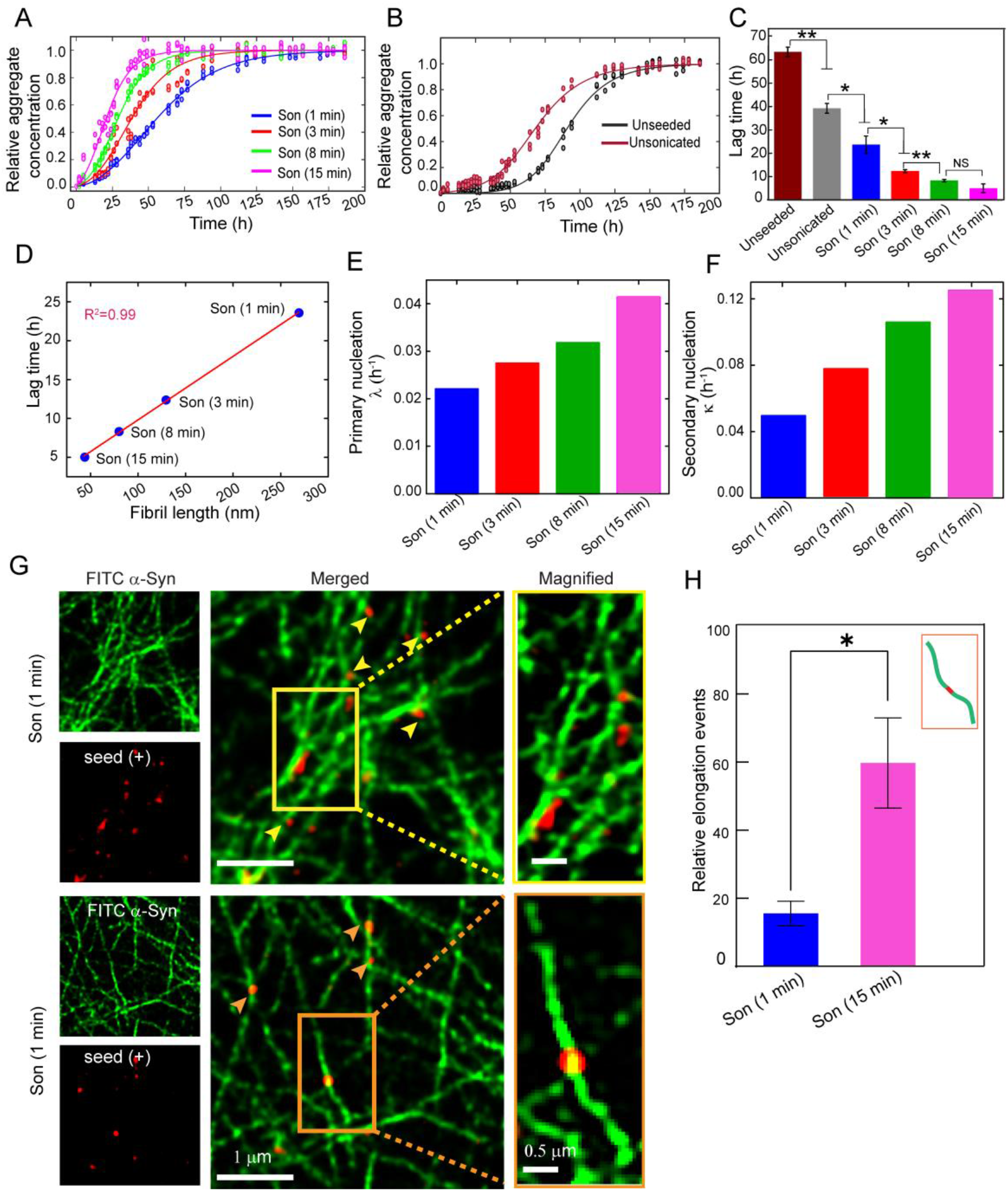
Size of α-Syn fibrils seeds dictate the microscopic steps involved in seeded amyloid assembly. (A) Aggregation kinetics of α-Syn (using ThT fluorescence) showing sigmoidal growth kinetics in the presence of 0.5% α-Syn fibrils seeds with varying lengths prepared by time-dependent sonication. The acquired data sets in triplicates were normalized using Amylofit online software. The normalized data sets (circles) for each group were well fitted (solid lines) to the model ‘Secondary Nucleation Dominated’ in AmyloFit. (B) Aggregation of α-Syn (150 μM monomeric protein) in the absence of fibril seeds (unseeded) and the presence of unsonicated fibrils was also recorded for comparison of lag time. (C) The bar diagram showing the lag time for the aggregation of α-Syn in the presence and absence of α-Syn fibrilar seeds of varying lengths; lag time is represented as Mean ± SEM from three independent sets of experiments. (D) Correlation plot of lag time and fibril length (R^2^= 0.99) calculated from TEM images showing that short fibril fragments are more potent in seeding the aggregation of α-Syn *in vitro*. (E, F) The overall parameters governing primary (λ) and secondary nucleation pathways (κ) were estimated from the rate constant values of primary nucleation (k_n_), elongation (k_+_), and k_2_ (surface-mediated secondary nucleation) from the fitting of sigmoidal amyloid growth curve with mean residual error, MRE= 0.0019. Since the observed elongation rate constant (k_+_) values are inversely related to the length of fibril seeds, the combined parameters calculated for primary nucleation (λ) and secondary nucleation (κ) subsequently showed an increasing trend with the extent of sonication. (G) Two-color STED super-resolution microscopy images showing the direct observation of the microscopic steps involved in secondary nucleation pathways. Regions of interest (ROIs) were magnified. The amyloid aggregation of FITC-labelled α-Syn monomer seeded with Rh-labelled long fibril seeds (Son (1 min)) showing the newer fibril growth from the side/surface of seeds corresponding to ‘surface-mediated secondary nucleation’ (yellow arrowheads). However, the reaction seeded with short fibrils (Son (15 min)) showed elongation events (marked with orange arrowheads). i.e., FITC-labelled daughter fibrils appear to be originating from both ends of Rh-labelled fibril seeds (bidirectional growth from parent seeds) – the characteristic feature of templated elongation. (H) Bar diagram showing the quantification of elongation events (daughter fibrils elongating from the ends of fibril seed) from STED images using Image J analysis showing relatively more elongation events for short fibrils compared to long fibrils seed. Experiments were repeated more than thrice. The data represented as Mean ± SEM.

To directly demonstrate the difference in secondary nucleation pathways in the fibril amplification by different-sized seeds, we used well-established two-colour super-resolution imaging studies (Pinotsi et al., 2014) using Stimulated Emission Depletion Microscopy (STED) (Klar et al., 2000, Vicidomini et al., 2018). We set up the aggregation of FITC-labelled α-Syn monomeric protein (9:1 unlabelled: FITC-labelled protein) seeded with Rh-labelled α-Syn fibril seeds of the short and the longest lengths. For the fibrillation in the presence of short seeds (Son (15 min)), we observed frequent events, wherein the FITC-labelled daughter fibrils were found to be originating from both ends of Rh-labelled parent fibril seeds (Figure 4G bottom panel). This is an indication of ‘elongation events’ in seeded amyloid assembly, wherein elongation of the seeds may dominate. Our observations are well corroborated with previous studies where fibril elongation is demonstrated as a bidirectional growth process from the α-Syn seeds as confirmed by super-resolution dSTORM imaging (Pinotsi et al., 2014). In contrast, in the case of seeded aggregation with longer seeds (Son (1 min)), we noticed frequent events resembling the surface-catalyzed secondary nucleation mechanism (FITC-labelled daughter fibril growth from the side/surface of the Rh-labelled longer fibril seeds) with few elongation events (Figure 4G top panel). This suggests that the monomeric α-Syn can attach to the catalytic surface of longer seeds by transient interaction, which may further accelerate the amyloid assembly by secondary nucleation pathways (Kumari et al., 2021). Therefore our findings provide the direct observation for the prevalence of seed size-dependent distinct microscopic mechanisms in secondary nucleation. In addition, the number of seeding events for each reaction was determined, suggesting the occurrence of abundant elongation events in aggregation reactions seeded with short fragments relative to their longer counterparts (Figure 4H).

### Size-dependent cellular apoptosis and membrane damage potential of α-Syn fibrils seeds

Apart from cellular uptake and seeding ability, the size of the fibrils may also affect the cytotoxic potential (Xue et al., 2010, Tarutani et al., 2016). This is because the fragmentation events lead to variability in fibril’s ends, which in turn modulate the interaction of fibril seeds with the cell membrane. In this context, we compared the cytotoxic potential of α-Syn fibril fragments of different lengths using MTT (4,5-Dimethylthiazol-2-yl)-2,5-diphenyltetrazolium bromide) reduction assay in SH-SY5Y cells. We observed a direct correlation with the size of fibrils, where there was a remarkable decrease in viability of cells when treated with fragmented α-Syn fibrils (Figure 5A). Cells treated with short fibril fragments (Son (15 min)) showed significantly less cell viability (∼49%) compared to longer α-Syn fibril fragments (Son (1 min); ∼64%). The extent of cellular apoptosis was also quantified by double staining of cells with Annexin-PI followed by flow cytometry analysis (Figure 5B). The cells treated with short fibrils fragments (Son (15 min)) showed an about two-fold increase in apoptotic cell death (∼ 55.8%) compared to longer fibril fragments (Son (1 min); ∼ 27%) and unsonicated fibrils (∼ 17.6%) (Figure 5C). These observations can be correlated (R^2^= 0.99) with the higher cellular uptake efficiency of short fibril seeds (Figure 5D), which eventually may cause enhanced apoptotic neurodegeneration due to the increased fibril loads in cells. We further quantified the apoptotic cell death in differentiated SH-SY5Y (SH-SY5Y (D)) cells. The overall cell death profile due to fibril treatment was found to be relatively high in differentiated cells; however, the observed trend in cell death due to treatment with fragmented fibril was comparable to that of undifferentiated SH-SY5Y cells (Figure S8). Hereafter, to understand the possible mechanism underlying enhanced cytotoxicity of the short fibril seeds, we quantified the membrane damage potential of fragmented α-Syn fibrils *in vitro* by examining the extent of dye release using carboxyfluorescein (CF) encapsulated synthetic liposomes. Short fibril fragments (Son (15 min)) showed ∼85% of CF release followed by Son (8 min) ∼72%, Son (3 min) ∼ 51%, and Son (1 min) ∼30%. However, the extent of dye release was considerably less (∼24%) for unsonicated fibrils (Figure 5 E). This suggests that short α-Syn fragments with more available fibril termini may interact more with dye encapsulated lipid membrane vesicles and therefore damage or create pores in the membrane, resulting in increased dye release from the vesicle into the solution. Further, correlation studies comparing percentage dye release caused by fragmented fibrils with its cytotoxic potential (R^2^=0.98) and cellular fibril uptake independently (R^2^=0.96) indicate that short fibril fragments with higher uptake efficiency caused extensive membrane damage and exerted high cytotoxicity (Figure 5 F). Taken together, our results demonstrate that the cytotoxic potential, membrane damage ability, and PD-associated cellular apoptosis are crucially dependent on the physical attributes of α-Syn amyloid fibrils despite having similar chemical identities.

**Figure 5:**
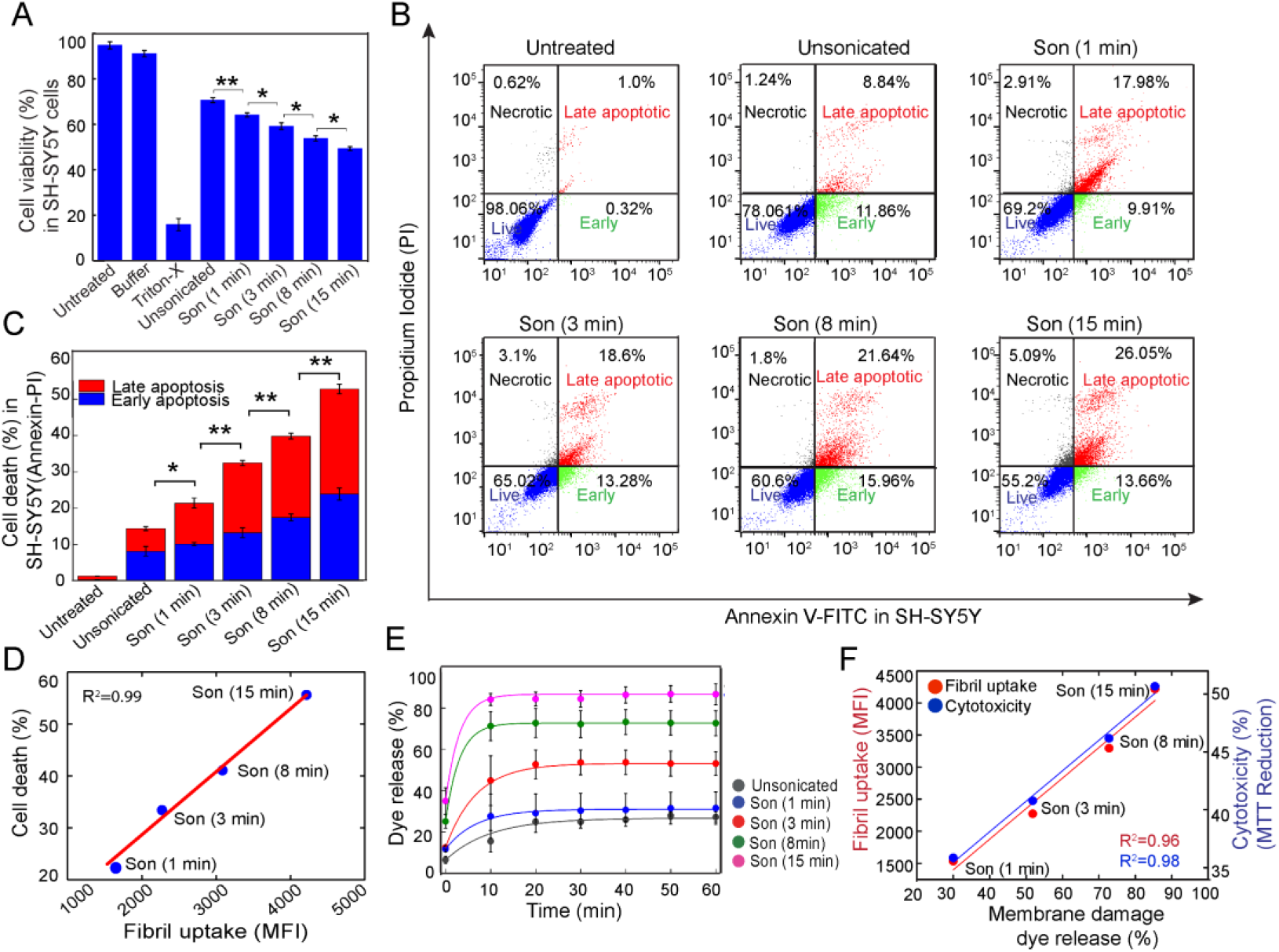
Size-dependent biological activities of α-Syn fibrils. (A) Cytotoxicity of α-Syn fibril fragments with variable lengths using MTT assay showing a decrease in cell viability with a decrease in the length of fibrils. Cells treated with unsonicated α-Syn fibrils were used as a control. (B) Flow cytometry analysis of cell death in Annexin V and PI stained cells showing the percentage of necrotic, late apoptotic, early apoptotic, and viable cell populations in SH-SY5Y cells upon treatment with α-Syn fibril fragments of varying lengths. (C) Bar plot showing an increase in apoptotic cell death in SH-SY5Ycells with a decrease in fibril length (for higher sonication time). The data represented as Mean ± SEM, calculated from three independent sets of experiments. (D) Correlation plot for the fibril uptake (MFI values from internalization data) with cell death quantified from the apoptotic assay in SH-SY5Y cells showing a positive correlation (R^2^= 0.99). (E) *In vitro* dye release assay indicating the membrane damage potential of α-Syn fibril fragments after binding to the CF-loaded synthetic liposomes. The data represented as Mean ± SEM analysed from three independent sets of experiments. (F) Correlation plot between membrane damage measured by dye release assay and cytotoxicity by MTT reduction (R^2^= 0.98) (blue) as well as fibril uptake acquired by FACS analysis (R^2^=0.96) (red).

## Discussion

An increasing body of evidence suggests that the prion-like behaviour of amyloid fibrils play a key role in the onset and spreading of various neurological disorders including PD (Karpowicz et al., 2019, Guo et al., 2016, Luk et al., 2012, Aguzzi and Lakkaraju, 2016, Eisenberg and Jucker, 2012). The prion-like propagation of amyloid fibrils and their associated cytotoxicity is thought to be determined by a series of events viz. fragmentation of fibrils and generation of seeds, followed by exponential amplification via amyloid growth pathways (Pezza and Serio, 2007, Xue et al., 2008, Xue et al., 2010, Marchante et al., 2017). For example, the infective potential of yeast prion protein Sup35NM depends on its physical dimensions, irrespective of the identical chemical nature of protein aggregates (Marchante et al., 2017).

The amyloid formation of α-Syn is a nucleation dependent polymerization reaction, which typically follows a sigmoidal growth pattern with three distinct phases (i) lag or nucleation phase, (ii) growth or elongation phase, and (iii) stationary or saturation phase (Wood et al., 1999, Ferrone, 1999, Knowles et al., 2014, Morris et al., 2009). However, the kinetic rate of this amyloid formation can be enhanced by the presence of PFF seeds, wherein the seed molecule hastens the amyloid transformation process (Harper and Lansbury, 1997, Arosio et al., 2015, Buell, 2017). Fibril fragmentation, leading to the formation of seeds, accelerates the amyloid assembly through secondary nucleation (Knowles et al., 2009, Xue et al., 2008, Xue et al., 2010, Gadhe et al., 2022). In this context, recent reports have shown critical evidence that secondary nucleation of amyloids also occurs in human cerebrospinal fluids (Frankel et al., 2019). Therefore, owing to the influence of PFF seeds, the secondary nucleation pathways have emerged as a central event in the autocatalytic amyloid amplification and its prion-like propagation (Meisl et al., 2020, Marrero-Winkens et al., 2020, Hadi Alijanvand et al., 2021). Molecular understanding of the seeded growth model suggests that secondary nucleation occurs either by templated elongation at the fibril termini (Pinotsi et al., 2014) or due to the catalytic nucleation pathways mediated by the fibril surfaces (Gaspar et al., 2017, Kumari et al., 2021). In the templated elongation pathway, the monomers are added at the growth competent ends of fibrils (Collins et al., 2004, Ferrone, 1999, Xue, 2015, Pinotsi et al., 2014), whereas surface-mediated growth involves catalysis of heterogeneous nucleation of new amyloid fibrils on the fibril surface (Gaspar et al., 2017, Zimmermann et al., 2021, Cohen et al., 2012, Kumari et al., 2021). These distinct microscopic steps affect the macroscopic protein aggregation processes abruptly, which may have significant implications in the intracellular transmission of various amyloids including α-Syn seeds in PD pathogenesis (Pinotsi et al., 2014, Gaspar et al., 2017, Linse, 2017, Mehra et al., 2021).

We hypothesize that the size of α-Syn fibril seeds might differentially affect the secondary nucleation pathways, during amyloid assembly. In the present study, we showed that controlled sonication can generate α-Syn amyloid fibrils of distinct fragment sizes without changing the major secondary structural features, cross-β-sheet conformation, or exposed hydrophobic surfaces (Figure 1). This allowed us to investigate how the size of α-Syn amyloids dictate the fibril amplification pathways. Since the uptake of fibrillar seeds is the initial step in the transcellular propagation of prion particles, we analysed the internalization behaviour of α-Syn fibril fragments in mammalian cells. Notably, we found a rapid and enhanced cellular uptake of the short fibril fragments compared to longer ones in human neuroblastoma and differentiated neuronal cells. This suggests that the cellular internalization behaviour of α-Syn fibril fragments strictly varies with seed length (Figure 2). These observations further suggest that the size-dependent complexity and the nanoscale differences in the physical attributes of amyloid fibrils could be sufficient to alter its cellular processes related to disease progression (Marchante et al., 2017, Beal et al., 2020). For instance; the binding of α-Syn PFF (preferably short fibril fragments) with its cell surface receptors could be more specific and selective (Mao et al., 2016), which consecutively facilitate its efficient cellular uptake and enhance the pathogenic inclusion formation in the mouse brain as compared to non-fragmented fibrils (Froula et al., 2019, Tarutani et al., 2016). Further, we showed that shorter fibrils have enhanced intracellular seeding potential/templating behaviour, which primarily drives the formation of elongated filamentous-like aggregates in the cytoplasm of the cells, overexpressing α-Syn. In contrast, small punctate-like intracellular protein aggregates were mostly generated by the exogenous addition of large fibril fragments in SH-SY5Y cells, overexpressing α-Syn (Figure 2 and Figure 3). Therefore, the difference in α-Syn seed size could dictate the distinct microscopic steps involved in the fibril amplification pathways in cells. This is well corroborated with previous studies demonstrating the mechanism of secondary nucleation involved in α-Syn aggregation *in vitro* using extensive structural analysis and super-resolution dSTORM imaging (Kumari et al., 2021, Pinotsi et al., 2014). Our observations further indicate that the generation and morphological features of these heterogeneous intracellular aggregates are largely dependent on fibril seed size. However, we speculate that these distinct morphological intracellular aggregates may further mature to form typical LB-like structures with time as reported previously (Mahul-Mellier et al., 2020).

Along these lines, we also tested the seed size-dependent secondary nucleation mechanism *in vitro* using global kinetic analysis together with super-resolution microscopy studies (Figure 4). All the fibril seeds irrespective of their sizes showed enhanced fibrillation kinetics of α-Syn; however, a much-reduced lag time was observed for the shortest fibril seeds. Our data suggest that the length of fibril seeds alters the overall rate of amyloid amplification possibly by modulating different secondary nucleation pathways such as seed-mediated elongation and seed-surface-mediated secondary nucleation. Super-resolution microscopy (using STED) studies of seeded aggregation indeed showed that short fibril seeds frequently accelerated the fibril formation via fibril elongation (by fibril termini) events, which eventually may engage in the fibril amplification process. In contrast, seeded aggregation with longer fibrils showed formation of fibrils from the surface of the fibril seeds. Here, it could be possible that longer fibril fragments with more surface area might induce the aggregation process preferably by surface-catalyzed secondary nucleation (Törnquist et al., 2018); possibly through the interaction of positively charged N-terminus of α-Syn monomer with C-terminus (negatively charged) of α-Syn fibrils seeds (Kumari et al., 2021) rather than by fibril elongation mechanism (Buell et al., 2014, Gaspar et al., 2017). Similarly, Knowles and co-workers have recently reported the demonstration of the surface-mediated secondary nucleation in seeded amyloid growth of Aβ using TIRF microscopy (Zimmermann et al., 2021). However, both the secondary nucleation processes are not exclusive of each other (Linse, 2019). They co-occur during the amplification process, but the extent of each of these processes varies with the seed length.

Apart from the differences in seed-size dependent amyloid amplification and internalization behaviour of α-Syn amyloid fibrils, we also observed a significant difference in their toxicity potential when exogenously added to undifferentiated and differentiated SH-SY5Y cells. Short fibril seeds consistently induced more cell death compared to longer fibrils; possibly due to the availability of more fibril termini leading to more interaction, at the cell membrane, for short seeds (as we used a similar concentration of seeds). Indeed, the *in vitro* assay of dye release from synthetic liposomes suggested that shorter fibril seeds caused more membrane damage, compared to longer fibrils seeds. Notably, the enhanced membrane damage ability and internalization behaviour of short fibril seeds accelerate the fibrils loads in cells, which might cause increased cytotoxicity and subsequent apoptotic neuronal cell death. (Figure 5). This is consistent with the recent *in vivo* study, wherein injection of the short α-Syn fibrillar fragments (∼29 nm in length) into the brain triggered extensive dopaminergic neurodegeneration, robust p-α-Syn inclusion formation, and motor dysfunction in a mouse model (Abdelmotilib et al., 2017) compared to the longer fibril fragments. This suggests that seed length of the amyloid fibril is a critical variable, which may control the neuropathological features such as onset, severity, and spreading of pathological phenotypes in PD.

In summary, our findings suggest that fragmentation of α-Syn fibrils generates amyloid fragments of varying physical dimensions with distinct internalization behaviour, seeding ability, and cytotoxic potential. The short fibril fragments with high uptake efficiency are the potent cytotoxic species, which accelerate amyloid amplification both *in vitro* and in cells. The nanoscale difference in the fragment length dictates distinct microscopic processes associated with the pathological conformational change of α-Syn. i.e., short fragments accelerate the amyloid growth by fibril elongation. In contrast, the longer fragments favour the surface-mediated secondary nucleation pathways for the amplification of α-Syn amyloid (Figure 6). Overall, this study helps to understand how the supra-structural variabilities, particularly the nanoscale differences in α-Syn fibrils seed length control the toxicity, amplification behaviour, prion-like transmission of α-Syn fibrils, and subsequent neurodegeneration in PD.

**Figure 6:**
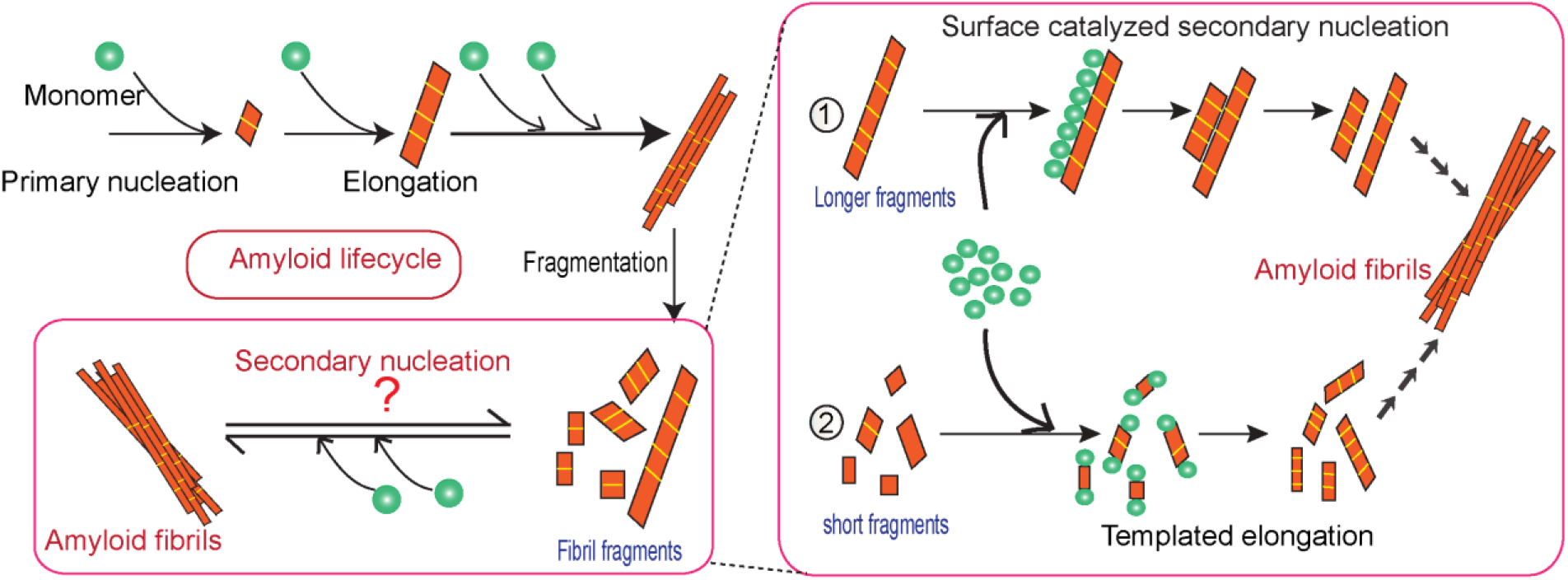
Scheme illustrating the impact of physical dimensions of fibril seeds on the mechanistic microscopic processes involved in the pathogenic conformational change of α-Syn. Aggregation of α-Syn occurs through a nucleation-dependent polymerization reaction. The molecular pathways underlying the structural transformation of α-Syn into its pathogenic conformation and its amplification rely on various microscopic steps majorly governed by amyloid fragmentation, and secondary nucleation. The physical dimensions of fibril seeds regulate the rate of these microscopic processes associated with secondary nucleation. For example, in an aggregation reaction seeded with longer fibril fragments; the extended fibril surface may act as a catalyst and mediate the surface-catalyzed secondary nucleation pathways ①. On the contrary, due to a higher number of growth competent fibril ends, the short fibril seeds template the aggregation by adding monomers on the fibril termini by elongation ②. Although these fibril seeds have similar chemical properties, their difference in physical dimensions alter their seeding potential and thus regulate the microscopic processes of amyloid assembly and the cellular processes associated with PD pathogenesis and progression.

## Methods

Expression and purification of α-Syn were carried out according to the established protocol (Volles and Lansbury, 2007, Singh et al., 2013, Ghosh et al., 2013) and used for fibrillation studies. α-Syn fibrils with varying seed lengths were rationally generated by a controlled time-dependent sonication procedure. The morphological characterization and size analysis of the fibril fragments was performed by TEM, AFM, and DLS studies. Biophysical characterization (including secondary structural features, exposed hydrophobic surfaces, and cross-β amyloid conformation) was studied by CD, FTIR, and XRD analysis. Moreover, the size-dependent effects of α-Syn fibril fragments in the cellular processes associated with disease onset/progression and cell death were evaluated in SH-SY5Y cells and differentiated neuronal cells using confocal microscopy and FACS analysis. The intracellular seeding potential and templating behavior of α-Syn fibril seeds with varying lengths were extensively studied in SH-SY5Y stable cells over-expressing C4-tagged α-Syn. The morphological studies and nature of the intracellular α-Syn aggregates were characterized by thorough imaging studies and aggresome staining assay (ProteoStat and flow cytometry). The seeding potential of fibril seeds with varying lengths and the resulting size-dependent fibril amplification pathways dictated by fibril seeds were analysed during the seeded aggregation kinetics of α-Syn monomer by ThT fluorescence assay. The global kinetic anlaysis of the data was fitted to a model ‘Secondary Nucleation Dominated’ using the Amylofit online platform (Meisl et al., 2016). The lag time and rate constant for primary nucleation (k_n_), elongation (k_+_), overall parameters governing primary nucleation (λ), secondary nucleation (κ) were calculated and compared among the groups. The distinct microscopic processes involved in secondary nucleation such as templated elongation and surface-mediated secondary nucleation dictated by fibril seeds with varying seeds were further demonstrated by dual-colour super-resolution STED microscopy and image analysis. The detailed methods are included in the supplementary information.

## Author contributions

The study was conceived and designed by S.K.M and A.S. All the experiments and data analysis were done by A.S. unless stated otherwise. D.D. carried out the data analysis of aggregation kinetics using Amylofit with the supervision of R.P. D.D. and K.P. performed confocal imaging and rendering of images. D.D. performed STED imaging and deconvolution of data with the guidance of K.S. A.N. performed FACS. L.G. helped in making dye-encapsulated lipid vesicles and protein expression. P.K. and R.K performed electron microscopy imaging. S.M. performed MALDI. D.C. helped in carrying out FTIR. A.S. and S.K.M wrote the manuscript. D.D., S.M., A.N. helped in editing the draft. All the authors have checked and approved the manuscript before submission.

## Acknowledgments

We are grateful to Dr. Juan Gerez and Prof. Roland Riek, ETH Zurich, Switzerland for the kind gift of the stable cell line, C4-tagged α-Syn SH-SY5Y. We thank Prof. Kamendra P. Sharma, Department of Chemistry, IIT Bombay for providing the DLS facility. We acknowledge IIT Bombay central facilities (TEM, FTIR, Bio-AFM, protein crystallography, MALDI-TOF/TOF mass spectrometer, confocal LSM/SD microscope) and the Department of Biosciences & Bioengineering for FACS facilities. We are thankful to the Microscopy facility at IISER Pune for performing super-resolution STED microscopy. We acknowledge DBT [BT/PR22749/BRB/10/1576/2016], Govt. of India for the funding. A.S. acknowledges ICMR-Research Associateship, Govt. of India (Sanction number: 2017/ 3466-CMB/BMS dated 02/04/2018) for fellowship.

## Supplementary information

Supplementary information includes detailed experimental methods, supplementary results (eight figures), and one table.

## References

1. Abdelmotilib, H., Maltbie, T., Delic, V., Liu, Z., Hu, X., Fraser, K. B., Moehle, M. S., Stoyka, L., Anabtawi, N., Krendelchtchikova, V., Volpicelli-Daley, L. A. & West, A. 2017. α-Synuclein fibril-induced inclusion spread in rats and mice correlates with dopaminergic Neurodegeneration. Neurobiol Dis, 105, 84–98.

2. Aguzzi, A. & Lakkaraju, A. K. 2016. Cell biology of prions and prionoids: a status report. Trends in cell biology, 26, 40–51.

3. Aguzzi, A. & Rajendran, L. 2009. The transcellular spread of cytosolic amyloids, prions, and prionoids. Neuron, 64, 783–90.

4. Arosio, P., Knowles, T. P. & Linse, S. 2015. On the lag phase in amyloid fibril formation. Phys Chem Chem Phys, 17, 7606–18.

5. Beal, D. M., Tournus, M., Marchante, R., Purton, T. J., Smith, D. P., Tuite, M. F., Doumic, M. & Xue, W. F. 2020. The Division of Amyloid Fibrils: Systematic Comparison of Fibril Fragmentation Stability by Linking Theory with Experiments. iScience, 23, 101512.

6. Buell, A. K. 2017. The nucleation of protein aggregates-from crystals to amyloid fibrils. International review of cell and molecular biology, 329, 187–226.

7. Buell, A. K., Galvagnion, C., Gaspar, R., Sparr, E., Vendruscolo, M., Knowles, T. P. J., Linse, S. & Dobson, C. M. 2014. Solution conditions determine the relative importance of nucleation and growth processes in α-synuclein aggregation. Proceedings of the National Academy of Sciences, 111, 7671–7676.

8. Chiti, F. & Dobson, C. M. 2006. Protein Misfolding, Functional Amyloid, and Human Disease. Annual Review of Biochemistry, 75, 333–366.

9. Cohen, S. I., Linse, S., Luheshi, L. M., Hellstrand, E., White, D. A., Rajah, L., Otzen, D. E., Vendruscolo, M., Dobson, C. M. & Knowles, T. P. 2013. Proliferation of amyloid-β42 aggregates occurs through a secondary nucleation mechanism. Proceedings of the National Academy of Sciences, 110, 9758–9763.

10. Cohen, S. I., Vendruscolo, M., Dobson, C. M. & Knowles, T. P. 2012. From macroscopic measurements to microscopic mechanisms of protein aggregation. Journal of molecular biology, 421, 160–171.

11. Collins, S. R., Douglass, A., Vale, R. D. & Weissman, J. S. 2004. Mechanism of prion propagation: amyloid growth occurs by monomer addition. PLoS Biol, 2, e321.

12. Dobson, C. M. 1999. Protein misfolding, evolution and disease. Trends in Biochemical Sciences, 24, 329–332.

13. Eisenberg, D. & Jucker, M. 2012. The amyloid state of proteins in human diseases. Cell, 148, 1188–1203.

14. Ferrone, F. 1999. Analysis of protein aggregation kinetics. Methods Enzymol, 309, 256–74.

15. Frankel, R., Törnquist, M., Meisl, G., Hansson, O., Andreasson, U., Zetterberg, H., Blennow, K., Frohm, B., Cedervall, T., Knowles, T. P. J., Leiding, T. & Linse, S. 2019. Autocatalytic amplification of Alzheimer-associated Aβ42 peptide aggregation in human cerebrospinal fluid. Communications Biology, 2, 365.

16. Frost, B., Jacks, R. L. & Diamond, M. I. 2009. Propagation of tau misfolding from the outside to the inside of a cell. The Journal of biological chemistry, 284, 12845–12852.

17. Froula, J. M., Castellana-Cruz, M., Anabtawi, N. M., Camino, J. D., Chen, S. W., Thrasher, D. R., Freire, J., Yazdi, A. A., Fleming, S., Dobson, C. M., Kumita, J. R., Cremades, N. & Volpicelli-Daley, L. A. 2019. Defining α-synuclein species responsible for Parkinson’s disease phenotypes in mice. The Journal of biological chemistry, 294, 10392–10406.

18. Gadhe, L., Sakunthala, A., Mukherjee, S., Gahlot, N., Bera, R., Sawner, A. S., Kadu, P. & Maji, S. K. 2022. Intermediates of α-synuclein aggregation: Implications in Parkinson’s disease pathogenesis. Biophysical Chemistry, 281, 106736.

19. Gaspar, R., Meisl, G., Buell, A. K., Young, L., Kaminski, C. F., Knowles, T. P. J., Sparr, E. & Linse, S. 2017. Secondary nucleation of monomers on fibril surface dominates α-synuclein aggregation and provides autocatalytic amyloid amplification. Q Rev Biophys, 50, e6.

20. Ghosh, D., Mondal, M., Mohite, G. M., Singh, P. K., Ranjan, P., Anoop, A., Ghosh, S., Jha, N. N., Kumar, A. & Maji, S. K. 2013. The Parkinson’s disease-associated H50Q mutation accelerates α-Synuclein aggregation in vitro. Biochemistry, 52, 6925–7.

21. Goedert, M., Eisenberg, D. S. & Crowther, R. A. 2017. Propagation of Tau Aggregates and Neurodegeneration. Annu Rev Neurosci, 40, 189–210.

22. Gómez-Benito, M., Granado, N., García-Sanz, P., Michel, A., Dumoulin, M. & Moratalla, R. 2020. Modeling Parkinson’s Disease With the Alpha-Synuclein Protein. Frontiers in Pharmacology, 11.

23. Guo, J. L., Narasimhan, S., Changolkar, L., He, Z., Stieber, A., Zhang, B., Gathagan, R. J., Iba, M., Mcbride, J. D., Trojanowski, J. Q. & Lee, V. M. 2016. Unique pathological tau conformers from Alzheimer’s brains transmit tau pathology in nontransgenic mice. J Exp Med, 213, 2635–2654.

24. Hadi Alijanvand, S., Peduzzo, A. & Buell, A. K. 2021. Secondary nucleation and the conservation of structural characteristics of amyloid fibril strains. Frontiers in Molecular Biosciences, 8, 268.

25. Hansen, C., Angot, E., Bergström, A. L., Steiner, J. A., Pieri, L., Paul, G., Outeiro, T. F., Melki, R., Kallunki, P., Fog, K., Li, J. Y. & Brundin, P. 2011. α-Synuclein propagates from mouse brain to grafted dopaminergic neurons and seeds aggregation in cultured human cells. J Clin Invest, 121, 715–25.

26. Harper, J. D. & Lansbury, P. T., Jr. 1997. Models of amyloid seeding in Alzheimer’s disease and scrapie: mechanistic truths and physiological consequences of the time-dependent solubility of amyloid proteins. Annu Rev Biochem, 66, 385–407.

27. Irtegun, S., Ramdzan, Y. M., Mulhern, T. D. & Hatters, D. M. 2011. ReAsH/FlAsH labeling and image analysis of tetracysteine sensor proteins in cells. J Vis Exp.

28. Jucker, M. & Walker, L. C. 2018. Propagation and spread of pathogenic protein assemblies in neurodegenerative diseases. Nature neuroscience, 21, 1341–1349.

29. Karpowicz, R. J. JR., Trojanowski, J. Q. & Lee, V. M. 2019. Transmission of α-synuclein seeds in neurodegenerative disease: recent developments. Lab Invest, 99, 971–981.

30. Klar, T. A., Jakobs, S., Dyba, M., Egner, A. & Hell, S. W. 2000. Fluorescence microscopy with diffraction resolution barrier broken by stimulated emission. Proceedings of the National Academy of Sciences, 97, 8206–8210.

31. Knowles, T. P., Vendruscolo, M. & Dobson, C. M. 2014. The amyloid state and its association with protein misfolding diseases. Nat Rev Mol Cell Biol, 15, 384–96.

32. Knowles, T. P. J., Waudby, C. A., Devlin, G. L., Cohen, S. I. A., Aguzzi, A., Vendruscolo, M., Terentjev, E. M., Welland, M. E. & Dobson, C. M. 2009. An Analytical Solution to the Kinetics of Breakable Filament Assembly. Science, 326, 1533–1537.

33. Koloteva-Levine, N., Aubrey, L. D., Marchante, R., Purton, T. J., Hiscock, J. R., Tuite, M. F. & Xue, W.-F. 2021. Amyloid particles facilitate surface-catalyzed cross-seeding by acting as promiscuous nanoparticles. Proceedings of the National Academy of Sciences of the United States of America, 118.

34. Kordower, J. H., Chu, Y., Hauser, R. A., Freeman, T. B. & Olanow, C. W. 2008. Lewy body–like pathology in long-term embryonic nigral transplants in Parkinson’s disease. Nature Medicine, 14, 504–506.

35. Krammer, C., Schätzl, H. M. & Vorberg, I. 2009. Prion-like propagation of cytosolic protein aggregates: insights from cell culture models. Prion, 3, 206–212.

36. Kumari, P., Ghosh, D., Vanas, A., Fleischmann, Y., Wiegand, T., Jeschke, G., Riek, R. & Eichmann, C. 2021. Structural insights into α-synuclein monomer–fibril interactions. Proceedings of the National Academy of Sciences, 118, e2012171118.

37. Kundel, F., Hong, L., Falcon, B., Mcewan, W. A., Michaels, T. C. T., Meisl, G., Esteras, N., Abramov, A. Y., Knowles, T. J. P., Goedert, M. & Klenerman, D. 2018. Measurement of Tau Filament Fragmentation Provides Insights into Prion-like Spreading. ACS chemical neuroscience, 9, 1276–1282.

38. Lansbury, P. T. 1999. Evolution of amyloid: What normal protein folding may tell us about fibrillogenesis and disease. Proceedings of the National Academy of Sciences, 96, 3342–3344.

39. Lashuel, H. A. 2020. Do Lewy bodies contain alpha-synuclein fibrils? and Does it matter? A brief history and critical analysis of recent reports. Neurobiol Dis, 141, 104876.

40. Linse, S. 2017. Monomer-dependent secondary nucleation in amyloid formation. Biophysical reviews, 9, 329–338.

41. Linse, S. 2019. Mechanism of amyloid protein aggregation and the role of inhibitors. Pure and Applied Chemistry, 91, 211–229.

42. Luk, K. C., Kehm, V., Carroll, J., Zhang, B., O’brien, P., Trojanowski, J. Q. & Lee, V. M. Y. 2012. Pathological α-synuclein transmission initiates Parkinson-like neurodegeneration in nontransgenic mice. Science (New York, N.Y.), 338, 949–953.

43. Luk, K. C., Song, C., O’brien, P., Stieber, A., Branch, J. R., Brunden, K. R., Trojanowski, J. Q. & Lee, V. M.-Y. 2009. Exogenous α-synuclein fibrils seed the formation of Lewy body-like intracellular inclusions in cultured cells. Proceedings of the National Academy of Sciences, 106, 20051–20056.

44. Mahul-Mellier, A. L., Burtscher, J., Maharjan, N., Weerens, L., Croisier, M., Kuttler, F., Leleu, M., Knott, G. W. & Lashuel, H. A. 2020. The process of Lewy body formation, rather than simply α-synuclein fibrillization, is one of the major drivers of neurodegeneration. Proc Natl Acad Sci U S A, 117, 4971–4982.

45. Mao, X., Ou, M. T., Karuppagounder, S. S., Kam, T.-I., Yin, X., Xiong, Y., Ge, P., Umanah, G. E., Brahmachari, S., Shin, J.-H., Kang, H. C., Zhang, J., Xu, J., Chen, R., Park, H., Andrabi, S. A., Kang, S. U., Gonçalves, R. A., Liang, Y., Zhang, S., Qi, C., Lam, S., Keiler, J. A., Tyson, J., Kim, D., Panicker, N., Yun, S. P., Workman, C. J., Vignali, D. A. A., Dawson, V. L., Ko, H. S. & Dawson, T. M. 2016. Pathological α-synuclein transmission initiated by binding lymphocyte-activation gene 3. Science (New York, N.Y.), 353, aah3374.

46. Marchante, R., Beal, D. M., Koloteva-Levine, N., Purton, T. J., Tuite, M. F. & Xue, W. F. 2017. The physical dimensions of amyloid aggregates control their infective potential as prion particles. Elife, 6.

47. Marrero-Winkens, C., Sankaran, C. & Schätzl, H. M. 2020. From Seeds to Fibrils and Back: Fragmentation as an Overlooked Step in the Propagation of Prions and Prion-Like Proteins. Biomolecules, 10, 1305.

48. Mehra, S., Ahlawat, S., Kumar, H., Singh, N., Navalkar, A., Patel, K., Kadu, P., Kumar, R., Jha, N. N., Udgaonkar, J. B., Agarwal, V. & Maji, S. K. 2020. α-Synuclein aggregation intermediates form fibril polymorphs with distinct prion-like properties. bioRxiv, 2020.05.03.074765.

49. Mehra, S., Gadhe, L., Bera, R., Sawner, A. S. & Maji, S. K. 2021. Structural and Functional Insights into α-Synuclein Fibril Polymorphism. Biomolecules, 11, 1419.

50. Meisl, G., Kirkegaard, J. B., Arosio, P., Michaels, T. C. T., Vendruscolo, M., Dobson, C. M., Linse, S. & Knowles, T. P. J. 2016. Molecular mechanisms of protein aggregation from global fitting of kinetic models. Nature Protocols, 11, 252–272.

51. Meisl, G., Knowles, T. P. & Klenerman, D. 2020. The molecular processes underpinning prion-like spreading and seed amplification in protein aggregation. Current opinion in neurobiology, 61, 58–64.

52. Morris, A. M., Watzky, M. A. & Finke, R. G. 2009. Protein aggregation kinetics, mechanism, and curve-fitting: a review of the literature. Biochim Biophys Acta, 1794, 375–97.

53. Nachman, E., Wentink, A. S., Madiona, K., Bousset, L., Katsinelos, T., Allinson, K., Kampinga, H., Mcewan, W. A., Jahn, T. R., Melki, R., Mogk, A., Bukau, B. & Nussbaum-Krammer, C. 2020. Disassembly of Tau fibrils by the human Hsp70 disaggregation machinery generates small seeding-competent species. J Biol Chem, 295, 9676–9690.

54. Pezza, J. A. & Serio, T. R. 2007. Prion propagation: the role of protein dynamics. Prion, 1, 36–43.

55. Pinotsi, D., Buell, A. K., Galvagnion, C., Dobson, C. M., Kaminski Schierle, G. S. & Kaminski, C. F. 2014. Direct Observation of Heterogeneous Amyloid Fibril Growth Kinetics via Two-Color Super-Resolution Microscopy. Nano Letters, 14, 339–345.

56. Ray, S., Singh, N., Kumar, R., Patel, K., Pandey, S., Datta, D., Mahato, J., Panigrahi, R., Navalkar, A., Mehra, S., Gadhe, L., Chatterjee, D., Sawner, A. S., Maiti, S., Bhatia, S., Gerez, J. A., Chowdhury, A., Kumar, A., Padinhateeri, R., Riek, R., Krishnamoorthy, G. & Maji, S. K. 2020. α-Synuclein aggregation nucleates through liquid-liquid phase separation. Nat Chem, 12, 705–716.

57. Roberti, M. J., Bertoncini, C. W., Klement, R., Jares-Erijman, E. A. & Jovin, T. M. 2007. Fluorescence imaging of amyloid formation in living cells by a functional, tetracysteine-tagged alpha-synuclein. Nat Methods, 4, 345–51.

58. Scheckel, C. & Aguzzi, A. 2018. Prions, prionoids and protein misfolding disorders. Nature Reviews Genetics, 19, 405–418.

59. Shen, D., Coleman, J., Chan, E., Nicholson, T. P., Dai, L., Sheppard, P. W. & Patton, W. F. 2011. Novel cell-and tissue-based assays for detecting misfolded and aggregated protein accumulation within aggresomes and inclusion bodies. Cell biochemistry and biophysics, 60, 173–185.

60. Shorter, J. & Lindquist, S. 2004. Hsp104 catalyzes formation and elimination of self-replicating Sup35 prion conformers. Science, 304, 1793–7.

61. Singh, P. K., Kotia, V., Ghosh, D., Mohite, G. M., Kumar, A. & Maji, S. K. 2013. Curcumin modulates α-synuclein aggregation and toxicity. ACS Chem Neurosci, 4, 393–407.

62. Spillantini, M. G. & Goedert, M. 2000. The alpha-synucleinopathies: Parkinson’s disease, dementia with Lewy bodies, and multiple system atrophy. Ann N Y Acad Sci, 920, 16–27.

63. Spillantini, M. G., Schmidt, M. L., Lee, V. M., Trojanowski, J. Q., Jakes, R. & Goedert, M. 1997. Alpha-synuclein in Lewy bodies. Nature, 388, 839–40.

64. Steiner, J. A., Angot, E. & Brundin, P. 2011. A deadly spread: cellular mechanisms of α-synuclein transfer. Cell Death & Differentiation, 18, 1425–1433.

65. Tanaka, M., Collins, S. R., Toyama, B. H. & Weissman, J. S. 2006. The physical basis of how prion conformations determine strain phenotypes. Nature, 442, 585–9.

66. Tarutani, A., Suzuki, G., Shimozawa, A., Nonaka, T., Akiyama, H., Hisanaga, S. & Hasegawa, M. 2016. The Effect of Fragmented Pathogenic α-Synuclein Seeds on Prion-like Propagation. J Biol Chem, 291, 18675–88.

67. Tittelmeier, J., Sandhof, C. A., Ries, H. M., Druffel-Augustin, S., Mogk, A., Bukau, B. & Nussbaum-Krammer, C. 2020. The HSP110/HSP70 disaggregation system generates spreading-competent toxic α-synuclein species. Embo j, 39, e103954.

68. Törnquist, M., Michaels, T. C., Sanagavarapu, K., Yang, X., Meisl, G., Cohen, S. I., Knowles, T. P. & Linse, S. 2018. Secondary nucleation in amyloid formation. Chemical Communications, 54, 8667–8684.

69. Vicidomini, G., Bianchini, P. & Diaspro, A. 2018. STED super-resolved microscopy. Nature Methods, 15, 173–182.

70. Volles, M. J. & Lansbury, P. T. JR. 2007. Relationships between the sequence of alpha-synuclein and its membrane affinity, fibrillization propensity, and yeast toxicity. J Mol Biol, 366, 1510–22.

71. Whitt, M. A. & Mire, C. E. 2011. Utilization of fluorescently-labeled tetracysteine-tagged proteins to study virus entry by live cell microscopy. Methods, 55, 127–36.

72. Winkler, J., Tyedmers, J., Bukau, B. & Mogk, A. 2012. Hsp70 targets Hsp100 chaperones to substrates for protein disaggregation and prion fragmentation. J Cell Biol, 198, 387–404.

73. Wood, S. J., Wypych, J., Steavenson, S., Louis, J. C., Citron, M. & Biere, A. L. 1999. alpha-synuclein fibrillogenesis is nucleation-dependent. Implications for the pathogenesis of Parkinson’s disease. J Biol Chem, 274, 19509–12.

74. Xue, W.-F., Hellewell, A. L., Hewitt, E. W. & Radford, S. E. 2010. Fibril fragmentation in amyloid assembly and cytotoxicity: when size matters. Prion, 4, 20–25.

75. Xue, W.-F., Homans, S. W. & Radford, S. E. 2008. Systematic analysis of nucleation-dependent polymerization reveals new insights into the mechanism of amyloid self-assembly. Proceedings of the National Academy of Sciences, 105, 8926–8931.

76. Xue, W. F. 2015. Nucleation: The Birth of a New Protein Phase. Biophys J, 109, 1999–2000.

77. Xue, W. F., Hellewell, A. L., Gosal, W. S., Homans, S. W., Hewitt, E. W. & Radford, S. E. 2009. Fibril fragmentation enhances amyloid cytotoxicity. J Biol Chem, 284, 34272–82.

78. Zimmermann, M. R., Bera, S. C., Meisl, G., Dasadhikari, S., Ghosh, S., Linse, S., Garai, K. & Knowles, T. P. J. 2021. Mechanism of Secondary Nucleation at the Single Fibril Level from Direct Observations of Aβ42 Aggregation. J Am Chem Soc, 143, 16621–16629.

