## Supplementary Information for "Size-dependent secondary nucleation and amplification of α-synuclein amyloid fibrils"

**Sections** **Pages**

1. Materials and Methods 2-18

2. Supplementary Results 19-22

3. Supplementary Figures (S1-S8) 23- 30

4. Supplementary Tables 31

5. Supplementary References 32-33

### Materials and Methods

#### Chemicals and reagents

Unless otherwise stated, high analytic grade chemicals obtained from Sigma Chemical Co. (St. Louis, MO, USA) were used for the study. Rhodamine dyes and FITC probes were purchased from Thermo Fisher Scientific. 1-palmitoyl-2-oleoyl-sn-glycero-3-phosphate (POP) and 1-palmitoyl-2-oleoyl-sn-glycero-3- phosphocholine (POPC) were purchased from Avanti Polar Lipids Inc. USA. The biarsenical dye, FlAsH-EDT2 was procured from Cayman Chemicals, USA. FITC-AnnexinV apoptosis detection kit was procured from BD Biosciences, USA. The ProteoStat aggresome detection assay kit was obtained from Enzo Life sciences, USA. Opti-MEM was purchased from Invitrogen, USA. High-quality Tet system-approved FBS was obtained from Takara Bio USA. Double distilled/ deionized water was obtained from the Milli-Q system (Millipore Corp., Bedford, MA) and autoclaved before use.

#### Cell culture

Human neuroblastoma (SH-SY5Y) cells were purchased from National Centre for Cell Science (NCCS) Pune. Cells were maintained in Dulbecco's modified Eagle's medium (DMEM) (Himedia) supplemented with Fetal Bovine Serum (10%) (Gibco) and 1% penicillin-streptomycin antibiotic cocktail (Himedia). SH-SY5Y cells were allowed to differentiate by treatment with a combination of retinoic acid ($10^{-7}$M) and 12-O-tetradecanoylphorbol-13-acetate ($1.6 \times 10^{-8}$ M) as per the previously established protocol (Påhlman et al., 1984, Påhlman et al., 1981). The neurite lengths of the differentiated cells were calculated using Image J and compared with the undifferentiated SH-SY5Y cells. SH-SY5Y cells stably expressing C4-tagged α-Syn (a kind gift from Dr. Juan Gerez and Prof.

Roland Riek, ETH Zurich) were grown in DMEM containing 10% Tet system approved FBS, 1% antibiotic cocktail, and used for intracellular seeding experiments. The overexpression of C4-tagged α-Syn was induced by treatment with doxycycline hyclate (DOX) (Sigma, USA) in SH-SY5Y cells as reported previously (Ray et al., 2020, Mehra et al., 2020, Gerez et al., 2019).

**Expression and purification of α-Syn**

Human α-Syn bacterial expression plasmid (pRK172) was used for the expression of the protein (Jakes et al., 1994). The proteins were expressed in *E. coli* strain, BL21 (DE3), and purified according to the protocol established by Volles *et al.* with marginal modifications (Volles and Lansbury, 2007, Singh et al., 2013, Ghosh et al., 2013). Then, the protein purity was assessed by SDS-PAGE and MALDI-TOF mass spectrometry analysis.

**Preparation of low molecular weight (LMW) α-Syn**

Low molecular weight α-Syn monomeric protein was used for all the fibrillation studies. Lyophilized protein (α-Syn) was dissolved in 20 mM Gly-NaOH buffer (pH 7.4) containing 0.01% sodium azide. LMW was prepared by a previously established protocol (Singh et al., 2013, Ghosh and Maji, 2015).

**Preparation of α-Syn amyloid fibril fragments**

Aggregation of α-Syn was carried out with a concentration of ~300 μM LMW in 20 mM Gly-NaOH buffer, (pH 7.4) containing 0.01% sodium azide. Sterile microcentrifuge tubes containing protein solutions were kept in Echo Therm model RT11 rotating mixture (Torrey Pines Scientific, USA) with slight agitation (50 rpm) and placed in a 37°C incubator. The structural transition during the aggregation was analysed by circular dichroism (CD) spectroscopy. The α-Syn amyloid fibril formation was often monitored by Thioflavin T (ThT) fluorescence assay and further confirmed by TEM analysis. Subsequently, after confirming the β-sheet structure by CD spectroscopy, the fibrils were pelleted down by ultracentrifugation (Beckman Coulter) at 40,000 rpm for 30 min. Then, the fibril concentration was determined by measuring the concentration of supernatant at 280 nm and deducting it from the original protein concentration, which had been set up for initial aggregation (300 μM). Later, the pelleted fibrils were resuspended in a suitable buffer according to the necessary fibril concentration required for each assay. Fibrils fragments with varying sizes were prepared by controlled fragmentation using a probe sonicator (Sonics & Materials, Inc) with pulse (3 sec ON; 1 sec OFF) amplitude (20%) for different time points viz. 1 min, 3 min, 8 min, and 15 min, and denoted as Son (1 min), Son (3 min), Son (8 min) and Son (15 min) respectively. Biophysical characterization of fragmented fibrils was performed soon after the sonication procedure. The morphological and size analyses were carried out by TEM, AFM, and DLS analysis.

**Circular dichroism (CD) spectroscopy**

Fragmented α-Syn fibril samples (7.5 μM) diluted in 20 mM Gly-NaOH buffer (pH 7.4) were taken in a 0.1 cm path-length quartz cell (Hellma, Forest Hills, NY). CD spectra were acquired using the JASCO-1500 instrument (wavelength range of 198-260 nm) at 25°C. Each spectrum was scanned thrice. Three independent experiments were performed for each sample.

**Thioflavin T (ThT) fluorescence assay**

ThT fluorescence assay was performed soon after the sonication of α-Syn fibrils. The fragmented fibril samples (7.5 μM) in 400 μL Gly-NaOH buffer were mixed with 4 μL of Thio T (1mM). Immediately after addition, ThT fluorescence assay was done using JASCO

(FP 8500) spectrofluorometer with excitation at 450 nm and emission in the range of 465−500 nm. The slit width for both excitation and emission was kept at 5 nm. Fluorescence emission at 480 nm was noted for ThT data analysis. Three sets of experiments in triplicates were done for each sample preparation.

**Transmission electron microscopy (TEM)**

Immediately after sonication, 10 μL of fragmented fibrils (~40 μM) were spotted on the formvar-coated copper grid procured from Electron Microscopy Sciences (Fort PA, Washington) and kept for 5 min. Then, the fibril sample was gently wiped off from the grid with a filter paper followed by sequential washing of the grid with autoclaved distilled water and staining with 1% (w/v) uranyl formate solution. The grids were subjected for imaging using a transmission electron microscope PHILIPS CM-200 (Amsterdam, Netherlands) with a magnification of 6600X at 200 kV. Multiple images (12 -15) were captured for each sample. The length distribution of fragmented fibrils was analyzed from acquired TEM images using ImageJ analysis (more than 260 fibril fragments were counted), which were plotted and fitted with appropriate peak functions using Origin Pro 8.0 (Origin Pro, USA).

**Atomic force electron microscopy (AFM)**

The morphological analysis of α-Syn fibril fragments prepared by time-dependent sonication was examined by AFM (Asylum Research, USA). To do so, 10 μM of the fibril fragments were spotted on a clean mica sheet and kept for ~10 min at room temperature followed by washing with autoclaved distilled water and desiccated under vacuum. Tapping mode AFM images were acquired using a silicon cantilever at a scan rate of 1.0 Hz.

**Dynamic light scattering (DLS)**

The particle size analysis of the fragmented fibrils prepared by time-dependent sonication was analysed by DLS. Freshly prepared fibril preparations (10 µM) were diluted in 1 mL filtered buffer (20 mM Gly-NaOH, pH 7.4). The particle measurements were taken in Anton Par Litesizer TM 500. The intensity distribution of particle size for each sample was plotted and compared. The measurement was recorded from 3 independent fibril preparations. The average particle size in terms of hydrodynamic diameter was determined by the inbuilt software based on the Brownian motion of the particles using the Stokes-Einstein equation as described previously (Hynes, 1977).

$dH = kT/3\pi\eta D$

Wherein, dH = hydrodynamic diameter, k = Boltzmann's constant, T= absolute temperature, $\eta$= viscosity and D = diffusion coefficient.

**Fourier Transform Infrared (FTIR) Spectroscopy**

For FTIR analysis, a small aliquot of fibril samples was spotted on a KBr pellet and permitted to dry with the help of an infrared lamp. The dried KBr pellet with the fibril samples was placed in a transmission holder in Bruker Vertex-80 instrument equipped with a DTGS detector. 32 scans were carried out for the amide I region (1500-1800 $cm^{-1}$). The frequency range (1700-1600 $cm^{-1}$) was deconvoluted by Fourier self-deconvolution method to examine the secondary structure (Kong and Yu, 2007). The deconvoluted spectra were subjected to curve fitting using Lorentzian function and were analysed using opus-65 software (Bruker, Leipzig, Germany) as per the manufacturer's recommendations. Multiple sets of FTIR spectra were recorded from three different fibril preparations.

**X-ray diffraction (XRD)**

For the X-ray diffraction study, high concentrations of the fragmented fibril samples were prepared and immediately transferred into a clean capillary tube (0.7 mm) and allowed to dry under vacuum. The capillary tube containing fibrils was fixed in the path of an X-ray beam for 200 sec at 1.2 kW. The diffraction patterns of the fibrils were recorded using the Rigaku R Axis IV++ detector (Rigaku, Japan) and examined using Adxv software. Three images were acquired for each group in duplicates.

**Sodium dodecyl sulfate-polyacrylamide gel electrophoresis (SDS-PAGE) analysis**

SDS-PAGE analysis was carried out to examine the purity of the α-Syn protein. Protein samples (25µM) were mixed with 5X SDS-gel loading buffer followed by heating for 10 min at 95°C in a water bath. Samples were then analysed on 15% SDS-PAGE and stained with Coomassie brilliant blue (CBB). The prestained protein ladder (Puregene Genetix, India) was also loaded on the same gel as a reference. Similarly, the stability of α-Syn fibril fragments prepared by time-dependent sonication was analysed by SDS-PAGE analysis. Soon after the sonication procedure, fibril fragments were analysed on 15% SDS-PAGE analysis followed by staining with CBB. Equal concentrations of unsonicated fibrils were also resolved on the same gel for comparison. Prestained full range protein ladder (Amersham Rainbow Marker, Merck) was used as a reference.

**Matrix-assisted laser desorption ionization-time of flight (MALDI-TOF) analysis of α-Syn**

α-Syn monomeric proteins were mixed with sinapinic acid (*trans*-3,5-dimethoxy-4-hydroxycinnamic acid, SA matrix) in a 1:1 ratio. Then, protein samples were spotted on a MALDI target plate and dried for 15-20 minutes at room temperature. MALDI-TOF mass spectrometric analysis was done using an Autoflex Speed (Bruker Daltonics) instrument. Data of crystallized spots were recorded by Flex control software followed by analysis of data as per the manufacturer's recommendation.

**NR binding Assay**

Immediately after sonication, the fibril fragments (10 µM) were diluted in buffer (20 mM Gly-NaOH, pH 7.4) and incubated for 5 min with 0.2 µl of 1 mM NR dye stock solution (prepared in DMSO). Fluorescence measurements were recorded with an excitation wavelength of 550 nm and 565-720 nm (emission wavelength range) using a spectrofluorometer (JASCO FP 8500). Three independent experiments in triplicates were performed.

**ANS binding assay**

To do ANS binding assay, 400 µl of diluted fibril fragments (10µM) in buffer (20 mM Gly-NaOH, pH 7.4) were incubated with 4µl of 5 mM ANS (8-anilino-1-naphthalenesulfonic acid) solution and allowed to incubate for 10 min in darkness. Then, fluorescence spectra were recorded using a spectrofluorometer (JASCO FP 8500) with an excitation wavelength of 370 nm and emission wavelength range from 400 nm to 600 nm. The excitation and emission slit width was kept at 5 nm. Three independent experiments in triplicates were performed. ANS control fluorescence was also measured with 4 µl ANS dye solution in the buffer.

**Aggregation Kinetics**

For studying the seeding ability of fragmented α-Syn fibrils with variable sizes, aggregation kinetics was performed with freshly prepared LMW (150 µM) in 20 mM Gly-NaOH buffer, (pH 7.4) containing 0.01% sodium azide in the presence of preformed α-Syn fibril fragments of variable lengths as seeds (0.5% v/v) at 37 $^{o}$C with slight agitation in 96-well clear bottom plate. 1.5 µL of ThT solution from 1 mM stock was added along with the samples to study the aggregation of the protein. Unseeded (LMW alone) and fibril seed alone were also incubated under the same conditions as controls. The ThT fluorescence signal was measured at regular intervals with excitation at 450 nm using a microplate reader (SpectraMax M2e, Molecular Devices, USA). The acquired data was fitted using AmyloFit online software (http://www.amylofit.ch.cam.ac.uk) following the guideline specified by Meisl et al. (Meisl et al., 2016). The best fit of the kinetics data was determined by using a basin hopping algorithm (Cohen et al., 2013). The relevant data range was chosen from the start of aggregation till a plateau was reached. The range was normalized by Amylofit and fitted according to the model 'Secondary Nucleation Dominated'.

The parameter of $k_n$ (primary nucleation rate constant), $k_+$ (elongation rate constant), $k_2$ (secondary nucleation rate constant) were estimated. κ is the overall parameter measuring the rate of secondary nucleation processes and λ is the parameter measuring the rate of primary nucleation processes (Knowles et al., 2009). The reaction order for both primary and secondary nucleations was considered as 'global constant' (n=2). $k_n$, primary nucleation rate constant was taken as a global fit, and $k_+$ and $k_2$ were taken as 'fit'. In this case, 10 basin hops were done to estimate each fit. The model estimates the best fit using the following equations (Meisl et al., 2016).

$$\frac{dP}{dt} = k_n m(t)^{nc} + k_2 m(t)^{n2} M(t)$$

$$\frac{dM}{dt} = 2m(t)k_+P(t)$$

$$\frac{M}{M_\infty} = 1 - \left(1 - \frac{M_0}{M_\infty}\right) e^{-k\infty t}$$

$$\left(\frac{B_- + C_+ e^{kt}}{B_+ + C_+ e^{kt}} \cdot \frac{B_+ + C_+}{B_- + C_+}\right)^{\frac{k\infty}{\kappa k\infty}}$$

The parameters were defined as,

$$\kappa = \sqrt{2m_0 k_+ m_0^{n2} k_2}$$

$$\lambda = \sqrt{2k_+ k_n m_0^{nc}}$$

$$C_\pm = \frac{k_+ P_0}{\kappa} \pm \frac{k_+ M_0}{2m_0 k_+} \pm \frac{\lambda^2}{2\kappa^2}$$

$$k_\infty = \sqrt{(2k_+ P(0))^2 + \frac{4k_+ k_n m_0^{nc}}{n_c} + \frac{4k_+ k_2 m_{tot} m_0^{n2}}{n_2} + \frac{4k_+ k_2 m_0^{n_2+1}}{n_2+1}}$$

$$\bar{k}_\infty = \sqrt{k_\infty^2 - 2C_+ C_- \kappa^2}$$

$$B_\pm = \frac{k_\infty \pm \bar{k}_\infty}{2\kappa}$$

The overall parameters, λ, and κ governing primary and secondary nucleation pathways were estimated from the values of $k_n$, $k_+$, and $k_2$ from the fitting.

The aggregation kinetics was repeated in three independent sets.

Then, the lag time for aggregation kinetics was determined using the following equation as reported previously (Willander et al., 2012).

y = y0 + (ymax – y0)/ (1+ e-(k(t-t1/2)),

Wherein, *y* indicates the ThT fluorescence at a particular time point,

*ymax* is the maximum ThT fluorescence and *y0* refers to the ThT fluorescence at *t0* and

*tlag* was defined as

t lag = t½- 2/k.

### Labelling of α-Synuclein

The commonly used amine-reactive fluorescent probes such as NHS-Rhodamine and FITC (Thermo fisher scientific, USA) were used for labelling the monomeric proteins. These probes attach and label at primary amines in the proteins moiety. To do so, monomeric protein solution (LMW) was prepared in non-amine containing conjugation buffers (PBS, pH 7.4 for Rhodamine and 0.1 M sodium bicarbonate, pH 9 for FITC), and labelling was done as per recommendations from manufacturer's guidelines (Thermo Fisher, Molecular Probes, USA). In brief, three molar excess of fluorescent probes (FITC and NHS-Rhodamine) were added slowly to the monomeric protein solution and incubated with constant stirring for 2 h at room temperature and another 6 h at 4 °C. Extensive dialysis (for ~48 h) in 1X PBS (pH 7.4) was carried out in a mini dialysis unit (Millipore, USA) against 20 mM Gly-NaOH, pH 7.4 with a frequent buffer exchange for every 3h to eliminate the unreacted dye from the protein solution. Then labelled protein was subjected to lyophilization and subsequently stored at $^{-}$20 $^{o}$C for further use.

**Preparation of Rhodamine (Rh)- labelled fibrils**

Labelled α-Syn monomer (2 mg) was dissolved in 200 µl of Gly- NaOH buffer (20 mM, pH 7.4). The degree of labelling and concentration of the proteins were determined according to the manufacturer's instructions. For fibrillation of Rh-labelled proteins, 90% of LMW α-Syn (unlabelled) was mixed with 10 % of Rh-labelled α-Syn at a protein concentration of 300 µM in 20 mM Gly-NaOH buffer containing 0.01% sodium azide (pH 7.4). This solution was kept in a rotational shaker (EchoTherm model RT11 (Torrey Pines Scientific, USA) at 50 rpm and incubated at 37 $^{o}$C and finally β-sheet conformation of labelled fibrils were confirmed by CD spectroscopy.

**Two-color super-resolution microscopy by STED**

Two-color STED imaging was performed to examine the microscopic process involved in seeded α-Syn amyloid formation. Since we need to correlate the size-dependent effect of α-Syn fibrils in secondary nucleation pathways, we compared seeds of two distinct sizes generated by time-dependent sonication. i.e., longer seeds (Son (1 min)) and short seeds (Son (15 min)). To visualize the microscopic events, the aggregation reaction was set up with 300 μM of FITC-labelled α-Syn (90% unlabelled LMW: 10% FITC-labelled LMW) in the presence of Rh-labelled α-Syn preformed fibrils as seeds as discussed above. The reaction mixture was kept in a rotational shaker (EchoTherm model RT11 (Torrey Pines Scientific, USA) at 37°C with slight agitation for 48h. Then samples were aliquoted and spotted on high-quality coverslips, the images were recorded in STED super-resolution microscopy (Leica Microsystems, Germany). The acquired images were further deconvoluted using Huygens software based on the CMLE algorithm.

**Internalization of fibrils in cells**

Concentration-dependent internalization experiments were performed in SH-SY5Y cells. 5 X $10^3$ cells per well were seeded on coverslips in a 12 well plate (Corning, USA) and incubated overnight at 37 °C with 5 % $CO_2$. Then cells were treated with increasing concentrations (50 nM, 100 nM, and 500 nM) of Rh-labelled α-Syn fibrils (fragmented/unfragmented) and incubated at 37 °C with 5 % $CO_2$ for 24h. The fragmented fibrils were prepared by standard sonication protocol for 3 minutes (with intermediate seed length) and used for the cellular uptake studies as established previously (Mehra et al., 2018). After 24 h, treated cells were washed twice with PBS (pH 7.4) to remove un-internalized fibrils and fixed with paraformaldehyde solution (4%) for 15 min. Then, fixed coverslips were mounted using mounting media containing DABCO (1, 4-Diazobicyclo-2,2,2-octane, Sigma) on a glass slide and observed under laser scanning confocal microscope (Carl Zeiss, LSM 780) with 63X oil immersion objective (1.4 NA). Similarly, the concentration-dependent cellular uptake of fibril fragments was further quantified by FACS analysis.

For studying size-dependent internalization behaviour of fibril fragments, SH-SY5Y cells (5 X $10^3$) per well were seeded on coverslips (18 mm) in a 12 well plate (Corning, USA) and incubated overnight at 37 °C with 5 % $CO_2$. Cells were then treated with 50 nM of Rh-labelled α-Syn fibrils fragments of variable length prepared by time-dependent sonication (1 min, 3 min, 8 min, 15 min) and incubated for 24 h. Similarly, cells treated with unsonicated fibrils were also used as control. After 24 h, treated cells were washed twice with PBS (pH 7.4) and fixed with paraformaldehyde solution (4%). Then, fixed coverslips were mounted on glass slides and observed under laser scanning confocal microscope (Carl Zeiss, LSM 780) with 63X oil immersion objective (1.4 NA). The imaging studies were performed thrice and image processing was carried out by Image J software.

The cellular uptake of fragmented fibrils with varying lengths was further quantified by FACS analysis. SH-SY5Y cells (3X $10^4$ cells) were seeded on a 24 well plate (Corning, USA) and grown at 37 °C with 5 % $CO_2$ overnight. Then, cells were treated with Rh-labelled fibrils of varying lengths (50 nM) for 24 h. Untreated and cells treated with buffer were used as controls. Cells were trypsinized after 24 h of incubation and analysed by FACS (BD Biosciences FACS Aria) in PE (561-A) channel. The absolute amount of fibrils internalized in each sample was calculated from respective Median Fluorescence Intensity (MFI) values. The gating was done with an untreated sample before analysis and 10,000 cells were recorded for each sample using the BD FACS Diva software. The recorded data was analysed and plotted using FlowJo v10 (Tree Star, Inc) software. For studying how incubation time affects the uptake behaviour of fragmented fibrils, time-dependent internalization experiments were performed by FACS analysis. SH-SY5Y cells (3X $10^4$ cells/well) were seeded on a 24 well plate and incubated overnight at 37 °C with 5 % $CO_2$. Then, cells were treated with 50 nM of fragmented α-Syn fibrils (Rh-labelled) of two distinct sizes (Son (1 min) and Son (15 min)) for two different time points of incubation (3h and 24h). Cells were collected by trypsinization after PBS wash and analysed by FACS. The gating was done with an untreated sample and 10,000 cells were recorded for each sample. Internalization experiments were performed at least 3 times for each condition.

**MTT assay of α-Syn fibril fragments**

To study the cytotoxic potential of α-Syn fibril fragments of variable size prepared by time-dependent sonication, MTT (3-(4, 5-dimethylthiazol2-yl)-2, 5 diphenyltetrazolium bromide) assay was performed in both SH-SY5Y and differentiated SH-SY5Y cells. Cells (1 X $10^4$ cells/well) were seeded in a 96 well plate (Corning, USA) and incubated for 24 h at 37$^o$C in 5% CO2. After that, cells were incubated with (25 μM) α-Syn fibril fragments of variable sizes prepared by time-dependent sonication. After 24 h, MTT reagent (0.5 mg/mL) was added per well and incubated for 4 h. Then solubilization buffer containing 50% DMF, 20% SDS (pH 4.7) was added and the plate was incubated overnight at 37$^o$C. After 16 hours, absorbance was taken at 560 nm and a background correction at 690 nm in a microplate reader (Molecular Devices, USA). Similarly, cells treated with buffer, unsonicated fibrils, and Triton-X were also analyzed. Three independent experiments were done in triplicates.

**Apoptosis detection assay**

SH-SY5Y cells (1X$10^5$ cells per well) were seeded in a 12-well culture plate (Corning, USA). After 24 h, cells were treated with 50 μM of α-Syn fibril fragments of varying lengths prepared by time-dependent sonication and incubated for 48 h. After the experimental period, cells were trypsinized and double-stained with Annexin V-FITC and PI according to the manufacturer's recommendations (Apoptosis Detection kit, BD Biosciences) and were quantified by flow cytometry (FACS Aria, BD Biosciences). 10,000 cells were counted for each sample and the recorded results were plotted using FlowJo v10 software (Tree Star, Inc). Cells treated with buffer and untreated cells were used as controls.

**Preparation of dye-loaded liposomes (SUVs)**

SUVs were prepared as established before (Ray et al., 2020). Briefly, Lipids viz. 1-palmitoyl-2-oleoyl-sn-glycero-3-phosphate (POP) and 1-palmitoyl-2-oleoyl-sn-glycero-3-phosphocholine (POPC) were mixed in a 1:3 molar ratio and dissolved in solvent (chloroform). The solvent was vacuum dried by evaporation by using a rotavap (Heidolph, Germany) at 50 rpm. After evaporation, a lipid layer was observed in the inner wall of the round bottom flask as a thin layer. This lipid film was then dissolved in sodium-phosphate buffer (20 mM, pH 7.4) containing 50 mM carboxyfluorescein (CF) by rotating the sample at 25 °C for at least 4 h in the dark. The lipid suspension was adjusted to a final concentration of 6 mM and the large multilamellar vesicles (LMVs) were sonicated to get SUVs by using a probe sonicator (Sonics and Materials Inc.; USA). The average size of the vesicles was ~100 nm as reported previously (Ray et al., 2020). The excess dye in the liposome preparation was removed by centrifugation for 30 min at 4℃ (18,000 g). The supernatant containing excess dye was discarded and the dye-loaded liposome pellet was again suspended in 20 mM sodium phosphate buffer (pH 7.4). This step was frequently repeated (at least thrice) to eliminate the excess of the dye.

**Membrane damage assay**

To analyse the membrane damage potential of α-Syn fibril fragments of variable size, dye leakage assay was done according to the previously established protocol (Singh et al., 2015) using CF-loaded liposomes. The CF-loaded liposomes were diluted 100 times with 20 mM sodium phosphate buffer (pH 7.4). α-Syn fibril fragments (10 μM) of variable lengths were prepared by time-dependent sonication and subsequently added to the diluted CF-loaded liposomes. Unsonicated fibrils were used as controls. Triton X-100 (0.5%) was kept as a positive control. The assay was performed in a clear bottom 96 well plate (Corning, USA) and fluorescence intensity (at 520 nm) was recorded (excitation at 495 nm) in a time-dependent manner at 25°C using a microplate reader (Molecular Devices, USA).

**In-cell seeding and quantification of intracellular aggregates**

SH-SY5Y cells stably expressing tetracysteine- tagged α-Syn (Gerez et al., 2019) were used for seeding assays. Cells (3 X $10^4$ per well) were seeded on 12 mm coverslips in a 24 well plate and maintained at 37 °C with 5 % CO2. The tetracysteine-tagged α-Syn expression was induced by incubating with doxycycline hyclate (DOX) for 24 h as reported previously (Ray et al., 2020, Mehra et al., 2020). Then, culture medium containing DOX was removed and treated for another 24 h with 50 nM of Rh-labelled fibril seeds of varying lengths. Then, the cells were stained with FlAsH-EDT2 using a previously established protocol (Ray et al., 2020, Roberti et al., 2007). In brief, cells were washed with PBS and incubated with 200 µL of 1 µM FlAsH and 10 mM EDT (dissolved in Opti-MEM) in a 37 °C with 5 % CO2 for 45 min. After that, cells were washed twice with PBS followed by 100 mM EDT to remove the unbound/ unreacted dye and then fixed the cells with ice-cold PFA (4%) for 5 min, then again washed with PBS pH 7.4. The fixed coverslips were immediately mounted on glass slides for imaging. Cell imaging was done under Zeiss Observer Z1 high-speed microlens-enhanced Nipkow spinning disc microscope (Zeiss, Germany) with 63X/1.4 NA oil immersion objective. Processing of the images and quantification of aggregates (red, green) and seeding events (yellow) were analysed by Image J software using a cell counter plugin. From these quantifications, we calculated the relative seeding event (%) by taking the ratio of observed seeding events (yellow) and internalized fibril seeds (red) per cell (Y/R). More than 80 cells were analyzed per sample from two independent sets of experiments. Further, the size analysis of the endogenous aggregates was also performed by image J and plotted the size distribution using MATLAB 2018b. 3D reconstruction of the cells with distinct morphological aggregates based on confocal Z-stack images was done by IMARIS 7.6.4 software (BitPlane, Switzerland).

**ProteoStat binding assay**

C4 α-Syn SH-SY5Y stable cells were used for the ProteoStat assay. Cells were seeded in 6 well plates and C4 α-Syn expression was induced in cells by treatment with DOX for 24h ($DOX^+$ samples) as reported previously (Mehra et al., 2020, Ray et al., 2020). 50 nM of α-Syn-fibril seeds of varying lengths were prepared by time-dependent sonication and incubated for 48 h. Similarly, uninduced (without DOX treated) cells were treated with 50 nM of α-Syn fibril samples ($DOX^-$ samples), used as respective controls. Cells were treated with a 5 μM aggresome inhibitor (MG132) and used as a positive control. After 48 h, cells were washed, fixed with 4% PFA, and permeabilized with 0.2% Triton X-100 solution followed by staining with 0.5μL of ProteoStat solution (5 μM in 5 ml of assay buffer) as per the manufacturer's instructions (Enzo Life Sciences, USA). Cells were subsequently analysed in the Texas Red channel by flow cytometry. The acquired data was analysed and plotted using FlowJo v10 (Tree Star, Inc) software. The aggresome propensity factor (APF) was further quantified from the mean fluorescence intensity (MFI) values as per the following formula.

$APF = 100 \times ((MFI_{DOX^+} - MFI_{DOX^-})/MFI_{DOX^+})$

Wherein, the mean fluorescence intensity (MFI) values obtained from $DOX^+$ cells treated with fibril seeds of varying lengths were denoted as $MFI_{DOX^+}$. Similarly, $MFI_{DOX^-}$ values for each group were used as control to deduct the fluorescent signal generated due to the binding of internalized α-Syn fibrils with the ProteoStat dye. MFI $_{DOX^-}$ values were acquired from $DOX^-$cells treated with α-Syn fibril seeds with respective seed lengths. The experiments were done three times.

**Statistical analysis**

The results are represented as Mean ± SEM from three independent sets of experiments. The statistical significance was analyzed by one-way ANOVA followed by Newman Keuls Multiple Comparison post hoc tests; the significant level is indicated as *P≤0.05; **P≤0.01; ***P≤0.001; **** P≤0.0001; NS (non-significant) P>0.05. KaleidaGraph, software was used for estimating the statistical significance among the groups.

### Supplementary Results

#### Generation of α-Syn fibril fragments and its characterization

We have expressed and purified recombinant α-Syn and the purity of the protein was validated by SDS-PAGE analysis and MALDI-TOF spectrometry (Figure S1 A, B). 300 μM of low molecular weight (LMW) protein was incubated at 37°C with slight agitation to prepare α-Syn fibrils. Meantime, the structural transition of protein was examined by CD spectroscopy. At the early stage of incubation, monomeric α-Syn showed the typical random coil (RC) structure and eventually converted to β-sheet rich fibrillar structure with single minima at ~218 nm (Figure S1C) at the end of the aggregation (after ~ 120 h). Subsequently, α-Syn fibrils were pelleted down by ultracentrifugation (40,000 rpm for 30 minutes) and resuspended in 20 mM Gly-NaOH buffer (pH 7.4). For generating α-Syn fibrils with variable lengths, controlled time-dependent sonication was performed using a probe sonicator (Sonics & Materials, Inc), which was used for further studies (Figure S1 D).

#### Optimization of fibril seed concentration for internalization experiments

It has been observed that cells treated with fragmented α-Syn fibrils prepared by standard sonication protocol for 3 minutes (with the intermediate length) showed fibril uptake in a concentration-dependent manner and formed distinct punctate-like structures in the cytoplasm. The unsonicated fibrils did not show any internalization and were observed in clumps adhering on the cell membrane (Figure S3 A). It may be due to the large size of the full-length fibrils, which may surpass the cut-off limit of what the cells can internalize. The cellular uptake of α-Syn fibrils was further quantified by FACS analysis. The overlaid histogram and corresponding median fluorescence intensity (MFI) values from FACS data also confirmed a concentration-dependent increase in the uptake of α-Syn fibrils fragments in cells (Figure S3 B, C); higher fibril uptake was recorded in cells treated with 500 nM fibrils (64,953 MFI) followed by 100 nM (20,354 MFI) and 50 nM (3027 MFI). Since α-Syn fragmented fibrils even at the lowest concentrations have shown significant cellular uptake, we have used this lowest concentration of α-Syn fibrils (50 nM) for our further internalization studies. In this case, the fluorescence signal saturation effect observed in higher fibril concentrations can also be neglected. Interestingly, α-Syn fibril seeds at lesser concentrations (50 nM) are observed to be nontoxic, which helped us to validate the size-dependent effects of fibril seeds with ease (Figure S7).

**The kinetics of the cellular uptake of short fibril seeds is higher compared to longer fibrils**

The size of fibril fragments not only affects the extent of internalization but also the exposure time required for its efficient cellular uptake. To examine this, time-dependent internalization experiments based on two different incubation periods (3 h and 24 h) were performed in SH-SY5Y cells to monitor the uptake pattern of α-Syn fibrils with two extreme sizes prepared by different time points of sonication such as Son (1 min) and Son (15 min). Cells treated with unsonicated fibrils were kept as control (Figure S3D). The time-dependent fibril uptake was quantitatively monitored by FACS analysis. Results indicate that cells treated with short fibril fragments (Son (15 min)) with an average length of ～44 nm showed a remarkably rapid uptake within 3 h and the extent of internalization of the short fibril seeds did not change significantly between 3 h (3671 MFI) and 24 h (3979 MFI) of incubation time. While cells treated with long fibril fragments (Son (1 min)) with an average length ～of 270 nm showed less internalization in 3 h of incubation (MFI 997); (i.e., exactly ～3.7 fold less fibril uptake as compared to cells treated with short fibril seeds for 3 h), which increased significantly (～1.6 fold) with an increase in incubation time (MFI 1627). It may be because the size of Son (1 min) fibril fragments are relatively long and hence may take a much longer incubation time (24 h) to show its highest internalization in cells. Moreover, the difference in fibril uptake was more observed in cells incubated with fibril fragments (short and long) for shorter periods (3h) as compared to extended (24h) incubation time (Figure S3D). Overall, these results suggest that the short α-Syn fibril fragments showed a very rapid internalization in SH-SY5Y cells relative to longer ones and the internalization of fibrils also depends on its exposure time.

**Differentiation of SH-SY5Y cells and cellular uptake of α-Syn fibril seeds**

Due to the technical challenge in using primary neuronal cells, SH-SY5Y neuroblastoma cells have been used widely for determining cellular toxicity of amyloid fibrils in various neurodegenerative disorders including PD (Xicoy et al., 2017). Further, to recapitulate the PD pathology, it has been widely accepted that SH-SY5Y neuroblastoma cells can be differentiated by treatment with various agents and thus making it a comparable cell model to mimic the disease phenotype (Presgraves et al., 2004, Lopes et al., 2010, Kovalevich and Langford, 2013)· Here, we have used a previously established differentiation protocol in which, treatment with retinoic acid followed by 12-O-tetradecanoylphorbol-13-acetate to get a neuronal phenotype (Påhlman et al., 1984). We observed significantly elongated neurite outgrowth in differentiated cells (Figure S5A, B). Next, we assessed the internalization behaviour of α-Syn fibril fragments in differentiated SH-SY5Y (SH-SY5Y (D)) cells. Identical treatment strategies and experimental conditions were used in (SH-SY5Y (D)) cells as used for undifferentiated cells. Similar to SH-SY5Y cells, a significant enhancement in the cellular uptake of the fragmented fibrils was observed, which is increased with decrease in fibril seed lengths. This was also confirmed using FACS analysis, where fibril internalization were quantified using Rh-labelled fibrils (Figure S5D, E). These observations were also well correlated with the corresponding confocal imaging data (Figure S5F). These observations collectively demonstrated that magnitude of fibril uptake in differentiated cells is consistent with the internalization pattern of fibril fragments observed in undifferentiated SH-SY5Y cells. In both cases, the internalization ability of α-Syn fibrils was dependent on its fibril seed length.

**Cytotoxicity of fibril seeds in differentiated SH-SY5Y cells**

The cytotoxic potential of fibril fragments of variable lengths was further evaluated in (SH-SY5Y(D)) cells using MTT assay (Figure S8B). The observed trend in results was consistent with the cell viability data observed in undifferentiated SH-SY5Y cells. We have also analyzed the apoptotic neuronal cell death upon treatment with 50 μM of α-Syn fibril fragments of variable lengths in SH-SY5Y (D) cells by flow cytometry (Figure S8C). Remarkably, we observed more cell death in differentiated cells relative to undifferentiated SH-SY5Y cells. It could be because the differentiated cells are more susceptible to neuronal toxicity exerted by amyloid fibril exposure. Moreover, the short fibril seeds caused extensive neuronal cell death relative to longer α-Syn seeds (Figure S8D) as observed for undifferentiated SH-SY5Y cells.

### Supplementary Figures

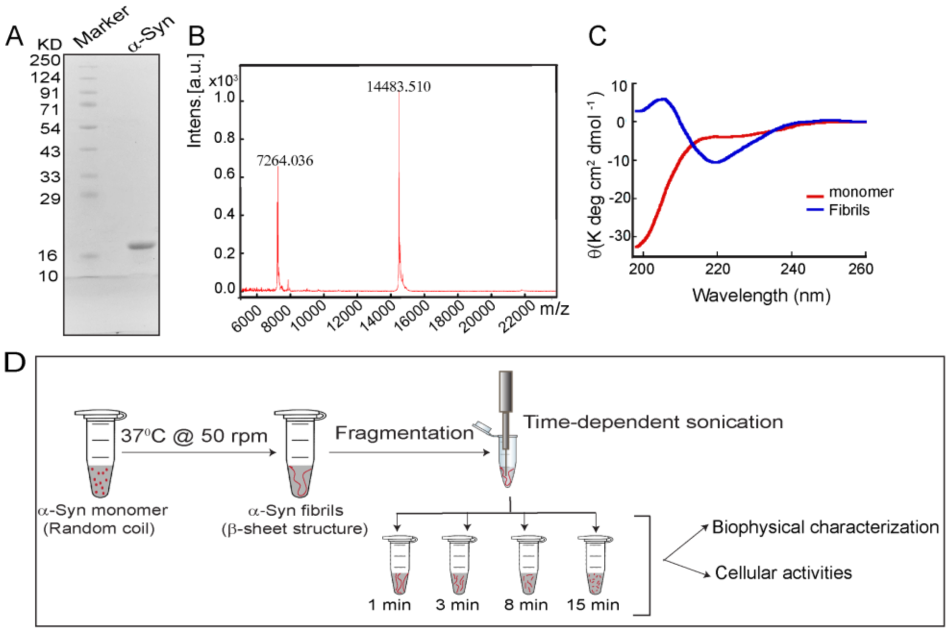


**Figure S1: Preparation of α-Syn fibrils fragments.** (A) 15% SDS-PAGE showing a single band at ~17 kDa corresponding to the purified α-Syn monomeric protein. (B) MALDI-TOF mass spectrometry indicates high intensity 14.4 kDa [M] and 7.2 kDa [M/2] peaks. (C) CD spectra showing the structural transition of the α-Syn from the random coil (RC) to β-sheet rich fibrillar conformation during α-Syn aggregation. (D) Scheme representing the generation of α-Syn fibril fragments of variable lengths by time-dependent sonication.

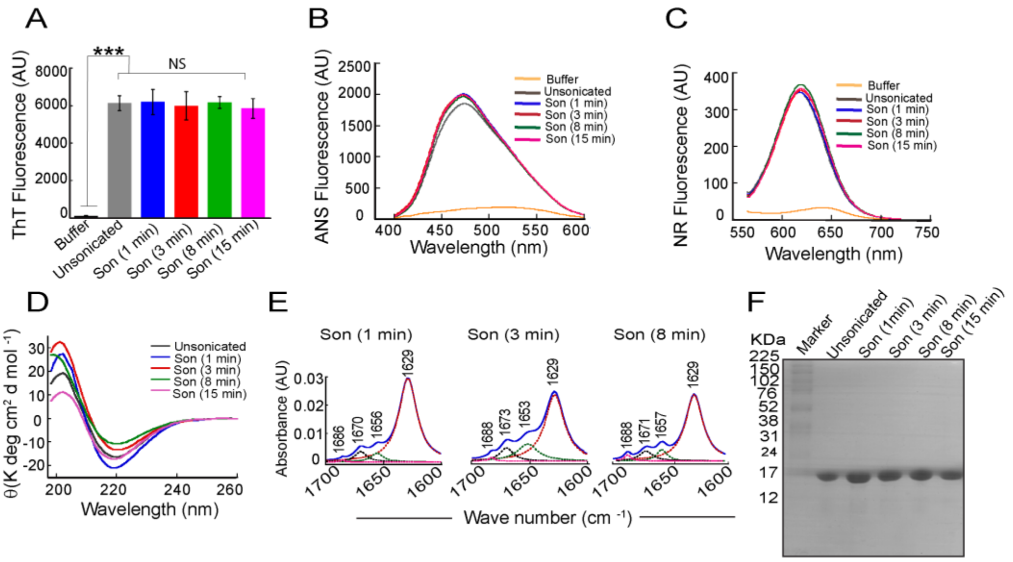


**Figure S2: Biophysical analysis of α-Syn fibril fragments.** (A) α-Syn fibrils with the variable size showing similar fluorescence intensity at 480 nm after binding with an amyloid-specific dye Thioflavin T. The values are represented as Mean ± SEM; n = 3 from independent experiments. (B) ANS fluorescence assay showing no alterations in the exposed hydrophobic surface and binding affinity of α-Syn fibrils with ANS after fragmentation. (C) NR fluorescence assay showing similar NR fluorescence after binding with different-sized fibrils seeds. (D) CD spectra of equimolar concentrations of α-Syn fibril fragments with the variable size showing β-sheet structure with a single minimum at ~218 nm; indicating no major changes in the secondary structure of α-Syn fibrils during the fragmentation events. The spectra were recorded for 3 independent sets of samples. (E) FTIR spectra of α-Syn fibrils prepared by time-dependent sonication showing, fragmentation events did not affect the secondary structure of α-Syn fibrils. (F) SDS-PAGE analysis of α-Syn fibrils fragments of variable size. After time-dependent sonication, fibril samples were analyzed using 15% SDS-PAGE. The single prominent band at ~17 kDa indicates no protein degradation occurred during the fragmentation procedure.

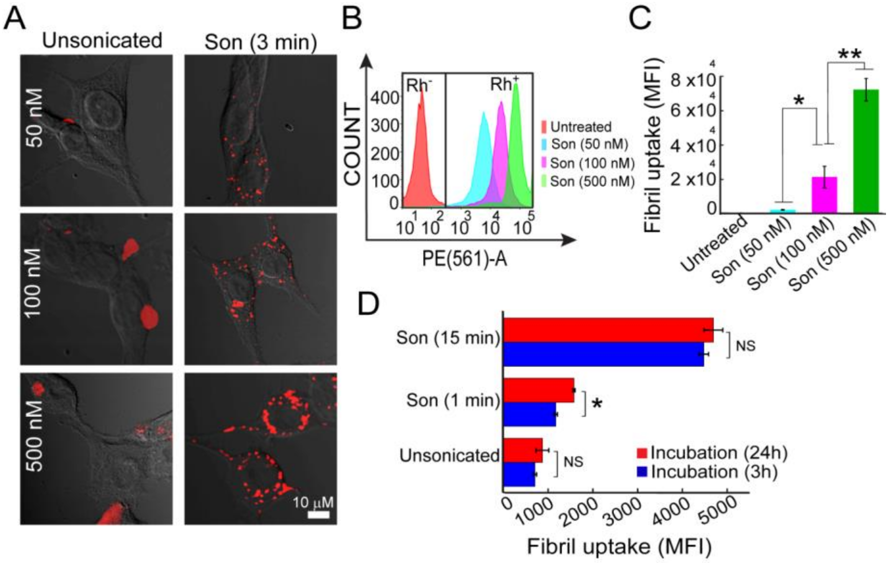


**Figure S3: Concentration-dependent studies for the internalization of α-Syn fibril fragments in SH-SY5Y cells.** (A) Confocal images of SH-SY5Y cells treated with Rhodamine (Rh)-labelled α-Syn fibril fragments of intermediate length (Son (3 min)) at three different fibril concentrations (50 nM, 100 nM, 500 nM) for 24 h showing concentration-dependent cellular uptake of the fragmented fibrils. The unsonicated fibrils did not get internalize and appeared as adhering to the cell membrane (confirmed by Z-stack imaging). The scale bar is 10 µm. (B) The overlaid histogram showing the concentration-dependent increase in $Rh^+$ cell population after internalization of α-Syn fibril fragments (Son (3 min) in SH-SY5Y cells. The gating for $Rh^+$ and $Rh^-$ was set based upon the fluorescence intensity of the untreated sample. (C) Bar plot showing the concentration-dependent significant difference in the median fluorescence intensity (MFI) of cells treated with α-Syn fibril fragments (50 nM, 100 nM, 500 nM). (D) Bar plot representing the quantification of internalized α-Syn fibril fragments suggesting that short fibril fragments (Son (15 min)) internalized very rapidly (3h) and showed saturated cellular uptake in 3h of incubation time; while, longer fibril fragments (Son (1 min)) showed increased internalization with time. Cells incubated with unsonicated fibrils for 3 h and 24 h were used as control.

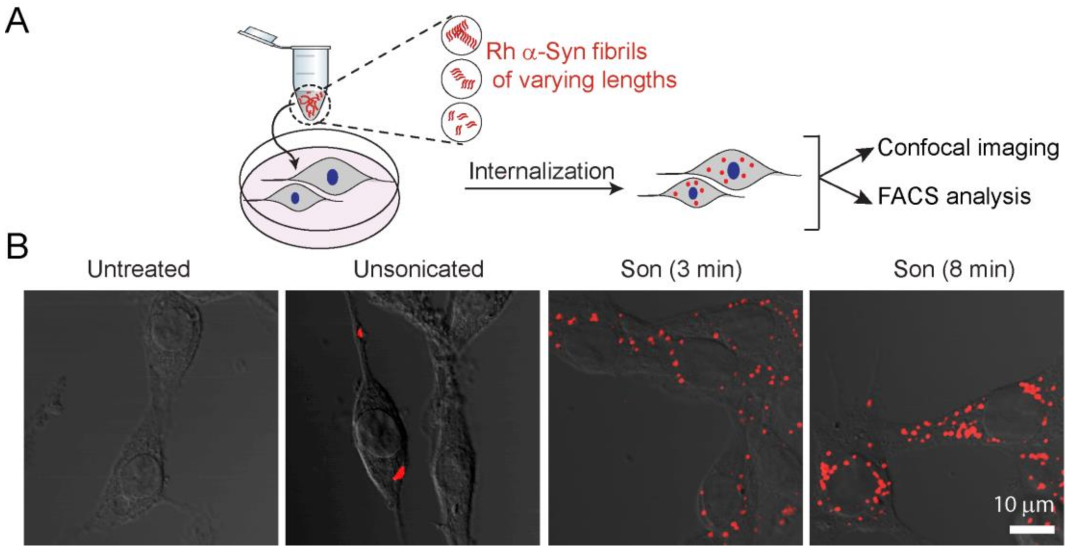


**Figure S4: Internalization of α-Syn fibril fragments with varying lengths in SH-SY5Y cells.** (A) Schematic illustration showing the experimental design for studying internalization of α-Syn fibril seeds with varying lengths and the punctate-formation in cells. (B) Confocal images showing fibril internalization by SH-SY5Y cells when treated for 24 h with Rh-labelled α-Syn fibril fragments (50 nM) of varying lengths (prepared by time-dependent sonication). Cells treated with unsonicated α-Syn fibrils were used as control.

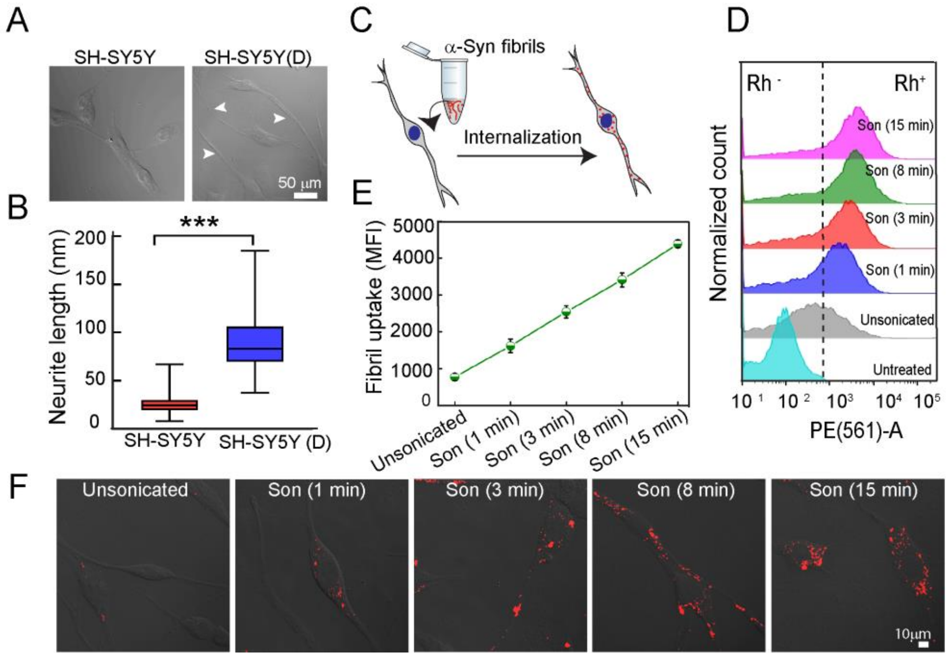


**Figure S5: Internalization behaviour of α-Syn fibril fragments with varying lengths in differentiated SH-SY5Y cells**. (A) Human neuroblastoma cells (SH-SY5Y, left panel) showing differentiation when treated with retinoic acid and phorbol ester (right panel). The differentiated SH-SY5Y (SH-SY5Y (D)) cells with extended neurites were indicated by white arrows. The scale bar is 50 µm. (B) Box plot showing the neurite length for differentiated and undifferentiated SH-SY5Y cells. (C) Schematic representation for the internalization studies in SH-SY5Y (D) cells. Rh-labelled α-Syn fibrils fragments were used for internalization in cells. Rh-labelled α-Syn fibrils fragments were internalized and formed punctate-like structures in cells. (D) The overlaid histogram showing the internalization efficiency of α-Syn fibril fragments (50 nM) with varying lengths in SH-SY5Y (D) cells. (E). Quantification for the cellular uptake (based on MFI values of cells acquired from FACS) showing an increase in the extent of internalization with an increase in sonication time. (F) Confocal images of differentiated cells showing fibril internalization after exogenous addition of different-sized Rh-labelled (50 nM) α-Syn fibril fragments. The sonication time used for generating different-sized fibril seeds was indicated in each image. Cells treated with unsonicated α-Syn fibrils were used as control. The scale bar is 10 µm.

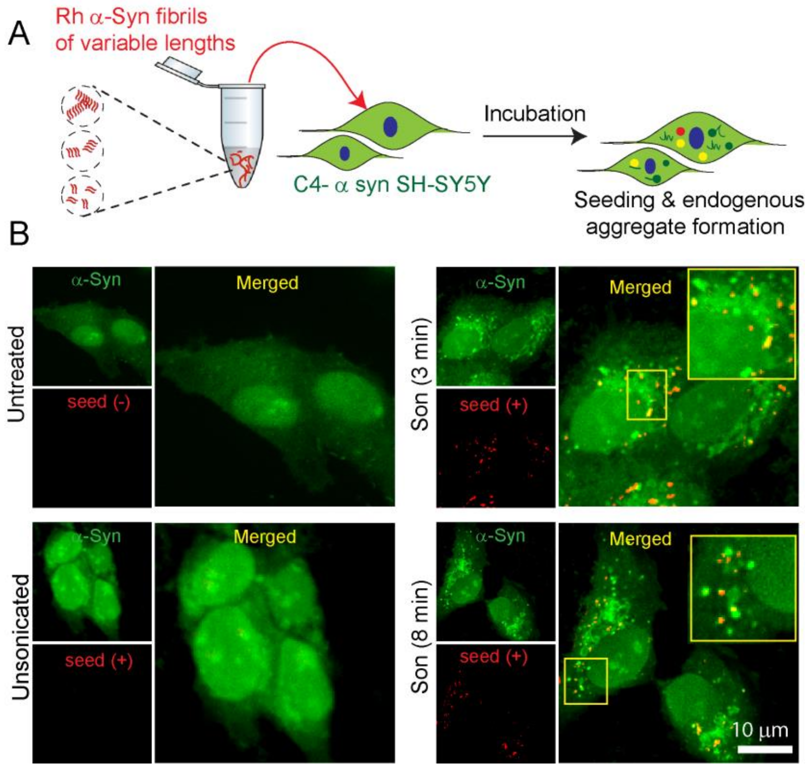


**Figure S6: Intracellular seeding potential of α-Syn fibril fragments with varying lengths.** (A) Schematic representation showing the experimental protocol used for studying the size-dependent seeding and amplification in cells. (B) Confocal images showing the seeding and amplification events for the endogenous aggregate formation observed in C4-tagged α-Syn SH-SY5Y cells after treatment with 50 nM of Rh-labelled α-Syn fibril seeds of variable lengths. Internalized α-Syn seeds were observed as red punctate structures. The merged images were magnified to highlight seeding events (colocalization, observed as yellow) and amplification of endogenous protein (observed as green inclusions) in cells. The intracellular seeding potential of longest (Son (1 min)) and shortest (Son (15 min)) α-Syn seeds was represented in Fig. 2D. The untreated cells were used as a control.

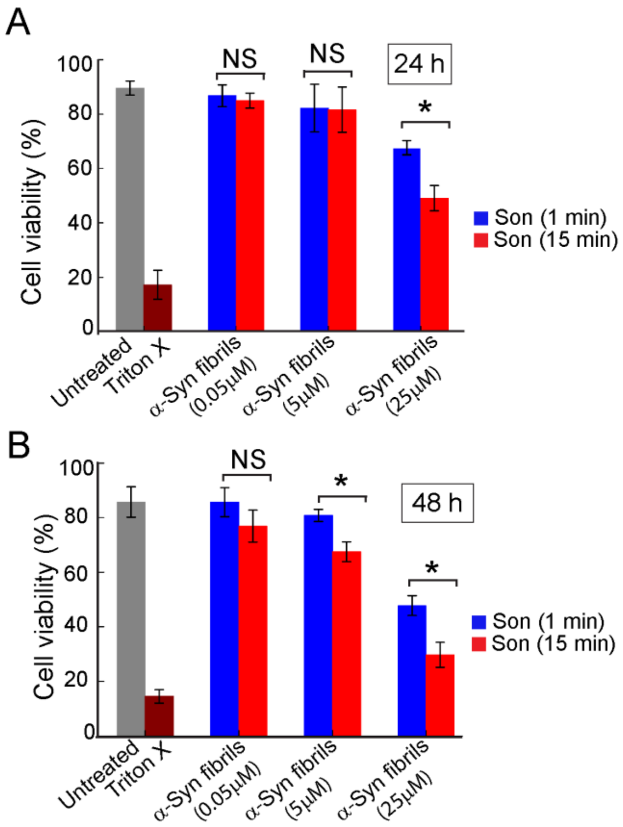


**Figure S7: Concentration-dependent toxicity assay for α-Syn fibril seeds of varying lengths in C4 α-Syn-tagged SH-SY5Y cells.** Bar plot indicating the percentage of cell viability examined in C4-tagged α-Syn overexpressing SH-SY5Y cells after treatment with increasing concentrations of α-Syn fibrils of two extreme seed lengths. The toxicity of the fibril seeds when incubated for 24 h (A) and 48h (B) was measured by MTT assay. Cells treated with Triton X-100 were used as a positive control.

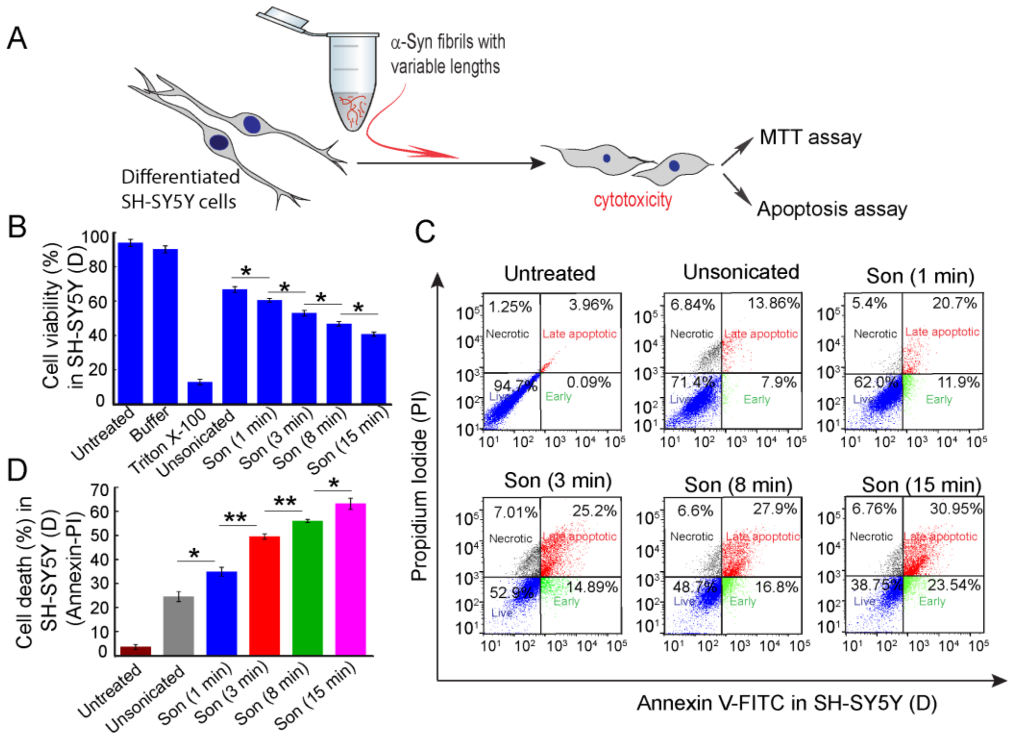


**Figure S8: Cytotoxic potential of α-Syn fibril seeds of varying lengths in differentiated SH-SY5Y (SH-SY5Y (D)) cells.** (A) Schematic representation showing the experimental procedure for toxicity assay in differentiated cells. (B) Cytotoxicity of α-Syn fibril fragments with variable lengths in SH-SY5Y(D) cells was quantified using MTT assay. Cells showed higher toxicity for shorter fibril fragments as compared to the longer ones. (C) FACS analysis using Annexin V and PI staining showing the percentage of necrotic, late apoptotic, early apoptotic, and viable populations in differentiated cells after treatment with different-sized fibrils. (D) The bar graph showing the total cell death in differentiated cells after treatment with different-sized fibril fragments.

**Table 1: Parameter values for estimating the rate constants for seed-dependent aggregation of α-Syn using Amylofit.**

| Seed Type (0.5%) | $m_0$ (M) | L (fibril length) | $M_0$ [ seed conc. (M)] | p (M/L) |
|---|---|---|---|---|
| **Son (1 min)** | 0.000150 | 270.46185 | 0.00000075 | 2.77303E-09 |
| **Son (3 min)** | 0.000150 | 129.4413 | 0.00000075 | 5.79413E-09 |
| **Son (8 min)** | 0.000150 | 80.019598 | 0.00000075 | 9.3727E-09 |
| **Son (15 min)** | 0.000150 | 43.898366 | 0.00000075 | 1.70849E-08 |

### References


1. COHEN, S. I., LINSE, S., LUHESHI, L. M., HELLSTRAND, E., WHITE, D. A., RAJAH, L., OTZEN, D. E., VENDRUSCOLO, M., DOBSON, C. M. & KNOWLES, T. P. 2013. Proliferation of amyloid-β42 aggregates occurs through a secondary nucleation mechanism. *Proceedings of the National Academy of Sciences,* 110**,** 9758-9763.
2. GEREZ, J. A., PRYMACZOK, N. C., ROCKENSTEIN, E., HERRMANN, U. S., SCHWARZ, P., ADAME, A., ENCHEV, R. I., COURTHEOUX, T., BOERSEMA, P. J., RIEK, R., PETER, M., AGUZZI, A., MASLIAH, E. & PICOTTI, P. 2019. A cullin-RING ubiquitin ligase targets exogenous α-synuclein and inhibits Lewy body–like pathology. *Science Translational Medicine,* 11**,** eaau6722.
3. GHOSH, D. & MAJI, S. K. 2015. Preparation of aggregate-free α-synuclein for in vitro aggregation study. *Protoc. Exchange***,** 270-281.
4. GHOSH, D., MONDAL, M., MOHITE, G. M., SINGH, P. K., RANJAN, P., ANOOP, A., GHOSH, S., JHA, N. N., KUMAR, A. & MAJI, S. K. 2013. The Parkinson's disease-associated H50Q mutation accelerates α-Synuclein aggregation in vitro. *Biochemistry,* 52**,** 6925-7.
5. HYNES, J. T. 1977. Statistical Mechanics of Molecular Motion in Dense Fluids. *Annual Review of Physical Chemistry,* 28**,** 301-321.
6. JAKES, R., SPILLANTINI, M. G. & GOEDERT, M. 1994. Identification of two distinct synucleins from human brain. *FEBS Lett,* 345**,** 27-32.
7. KNOWLES, T. P. J., WAUDBY, C. A., DEVLIN, G. L., COHEN, S. I. A., AGUZZI, A., VENDRUSCOLO, M., TERENTJEV, E. M., WELLAND, M. E. & DOBSON, C. M. 2009. An Analytical Solution to the Kinetics of Breakable Filament Assembly. *Science,* 326**,** 1533-1537.
8. KONG, J. & YU, S. 2007. Fourier transform infrared spectroscopic analysis of protein secondary structures. *Acta Biochim Biophys Sin (Shanghai),* 39**,** 549-59.
9. KOVALEVICH, J. & LANGFORD, D. 2013. Considerations for the use of SH-SY5Y neuroblastoma cells in neurobiology. *Methods Mol Biol,* 1078**,** 9-21.
10. LOPES, F. M., SCHRÖDER, R., DA FROTA, M. L., JR., ZANOTTO-FILHO, A., MÜLLER, C. B., PIRES, A. S., MEURER, R. T., COLPO, G. D., GELAIN, D. P., KAPCZINSKI, F., MOREIRA, J. C., FERNANDES MDA, C. & KLAMT, F. 2010. Comparison between proliferative and neuron-like SH-SY5Y cells as an in vitro model for Parkinson disease studies. *Brain Res,* 1337**,** 85-94.
11. MEHRA, S., AHLAWAT, S., KUMAR, H., SINGH, N., NAVALKAR, A., PATEL, K., KADU, P., KUMAR, R., JHA, N. N., UDGAONKAR, J. B., AGARWAL, V. & MAJI, S. K. 2020. α-Synuclein aggregation intermediates form fibril polymorphs with distinct prion-like properties. *bioRxiv***,** 2020.05.03.074765.
12. MEHRA, S., GHOSH, D., KUMAR, R., MONDAL, M., GADHE, L. G., DAS, S., ANOOP, A., JHA, N. N., JACOB, R. S., CHATTERJEE, D., RAY, S., SINGH, N., KUMAR, A. & MAJI, S. K. 2018. Glycosaminoglycans have variable effects on α-synuclein aggregation and differentially affect the activities of the resulting amyloid fibrils. *J Biol Chem,* 293**,** 12975-12991.
13. MEISL, G., KIRKEGAARD, J. B., AROSIO, P., MICHAELS, T. C. T., VENDRUSCOLO, M., DOBSON, C. M., LINSE, S. & KNOWLES, T. P. J. 2016. Molecular mechanisms of protein aggregation from global fitting of kinetic models. *Nature Protocols,* 11**,** 252-272.
14. PÅHLMAN, S., ODELSTAD, L., LARSSON, E., GROTTE, G. & NILSSON, K. 1981. Phenotypic changes of human neuroblastoma cells in culture induced by 12-O-tetradecanoyl-phorbol-13-acetate. *Int J Cancer,* 28**,** 583-9.
15. PÅHLMAN, S., RUUSALA, A.-I., ABRAHAMSSON, L., MATTSSON, M. E. K. & ESSCHER, T. 1984. Retinoic acid-induced differentiation of cultured human neuroblastoma cells: a comparison with phorbolester-induced differentiation. *Cell Differentiation,* 14**,** 135-144.

16. PRESGRAVES, S. P., AHMED, T., BORWEGE, S. & JOYCE, J. N. 2004. Terminally differentiated SH-SY5Y cells provide a model system for studying neuroprotective effects of dopamine agonists. *Neurotox Res,* 5**,** 579-98.
17. RAY, S., SINGH, N., KUMAR, R., PATEL, K., PANDEY, S., DATTA, D., MAHATO, J., PANIGRAHI, R., NAVALKAR, A., MEHRA, S., GADHE, L., CHATTERJEE, D., SAWNER, A. S., MAITI, S., BHATIA, S., GEREZ, J. A., CHOWDHURY, A., KUMAR, A., PADINHATEERI, R., RIEK, R., KRISHNAMOORTHY, G. & MAJI, S. K. 2020. α-Synuclein aggregation nucleates through liquid-liquid phase separation. *Nat Chem,* 12**,** 705-716.
18. ROBERTI, M. J., BERTONCINI, C. W., KLEMENT, R., JARES-ERIJMAN, E. A. & JOVIN, T. M. 2007. Fluorescence imaging of amyloid formation in living cells by a functional, tetracysteine-tagged alpha-synuclein. *Nat Methods,* 4**,** 345-51.
19. SINGH, P. K., GHOSH, D., TEWARI, D., MOHITE, G. M., CARVALHO, E., JHA, N. N., JACOB, R. S., SAHAY, S., BANERJEE, R., BERA, A. K. & MAJI, S. K. 2015. Cytotoxic helix-rich oligomer formation by melittin and pancreatic polypeptide. *PLoS One,* 10**,** e0120346.
20. SINGH, P. K., KOTIA, V., GHOSH, D., MOHITE, G. M., KUMAR, A. & MAJI, S. K. 2013. Curcumin modulates α-synuclein aggregation and toxicity. *ACS Chem Neurosci,* 4**,** 393-407.
21. VOLLES, M. J. & LANSBURY, P. T., JR. 2007. Relationships between the sequence of alpha-synuclein and its membrane affinity, fibrillization propensity, and yeast toxicity. *J Mol Biol,* 366**,** 1510-22.
22. WILLANDER, H., PRESTO, J., ASKARIEH, G., BIVERSTÅL, H., FROHM, B., KNIGHT, S. D., JOHANSSON, J. & LINSE, S. 2012. BRICHOS domains efficiently delay fibrillation of amyloid β-peptide. *Journal of Biological Chemistry,* 287**,** 31608-31617.
23. XICOY, H., WIERINGA, B. & MARTENS, G. J. M. 2017. The SH-SY5Y cell line in Parkinson's disease research: a systematic review. *Molecular Neurodegeneration,* 12**,** 10.
